# Meta-analytic clustering dissociates brain activity and behavior profiles across reward processing paradigms

**DOI:** 10.1101/818948

**Authors:** Jessica S. Flannery, Michael C. Riedel, Katherine L. Bottenhorn, Ranjita Poudel, Taylor Salo, Lauren D. Hill-Bowen, Angela R. Laird, Matthew T. Sutherland

**Author notes:** **Author Note:** This manuscript’s content is original research and is not currently in submission elsewhere while under consideration at *Cognitive, Affective, and Behavioral Neuroscience.* All authors have contributed substantially to this manuscript. There are no financial or other relations associated with this study that represent a conflict of interest. Correspondence: Matthew T. Sutherland, Ph.D. Florida International University Department of Psychology AHC-4, RM 312 11299 S.W. 8th St Miami, FL 33199.

## Abstract

Reward learning is a ubiquitous cognitive mechanism guiding adaptive choices and behaviors, and when impaired, can lead to considerable mental health consequences. Reward-related functional neuroimaging studies have begun to implicate networks of brain regions essential for processing various peripheral influences (e.g., risk, subjective preference, delay, social context) involved in the multifaceted reward processing construct. To provide a more complete neurocognitive perspective on reward processing that synthesizes findings across the literature while also appreciating these peripheral influences, we utilized emerging meta-analytic techniques to elucidate brain regions, and in turn networks, consistently engaged in distinct aspects of reward processing. Using a data-driven, meta-analytic, *k*-means clustering approach, we dissociated seven meta-analytic groupings (MAGs) of neuroimaging results (i.e., brain activity maps) from 749 experimental contrasts across 176 reward processing studies involving 13,358 healthy participants. We then performed an exploratory functional decoding approach to gain insight into the putative functions associated with each MAG. We identified a seven-MAG clustering solution which represented dissociable patterns of convergent brain activity across reward processing tasks. Additionally, our functional decoding analyses revealed that each of these MAGs mapped onto discrete behavior profiles that suggested specialized roles in predicting value (MAG-1 & MAG-2) and processing a variety of emotional (MAG-3), external (MAG-4 & MAG-5), and internal (MAG-6 & MAG-7) influences across reward processing paradigms. These findings support and extend aspects of well-accepted reward learning theories and highlight large-scale brain network activity associated with distinct aspects of reward processing.

## INTRODUCTION

Reward learning is a fundamental cognitive process vital for adaptive functioning that consequently, when disrupted, has a significant impact on mental health. Widely accepted and commonly adopted neurocognitive models of reward learning propose that value predictions are formed and updated based on dopaminergic reward prediction error (RPE) signals (Glimcher, 2011; Hollerman & Schultz, 1998; Knutson et al., 2001; Schultz, 1998; Schultz, 2016; Schultz, Dayan & Montague, 1997). RPE signals represent the discrepancy between expected outcomes and actual outcomes (Rescorla, 1972). In support of these models, functional neuroimaging studies have consistently demonstrated striatal and mid-brain activity associated with unexpected positive or negative cues/outcomes (i.e., RPE task events) across a variety of paradigms (Cooper et al., 2009; Garrison, Erdeniz & Done, 2013; Knutson et al., 2008; O’Doherty et al., 2003). As neuroimaging investigations of value prediction have advanced, scientists continue to construct progressively more complex paradigms involving reward contingencies within increasingly externally valid contexts involving additional influences on valuation, and ultimately reward learning (Daw et al., 2011; Kahnt, 2018; Porcelli & Delgado, 2009; Seaman et al., 2018).

For example, many reward processing tasks involve paradigms that mirror gambling activities played regularly in casinos or watched on game shows, like blackjack (Miedl et al., 2010), wheel of fortune (Ernst et al., 2005), and slot machines (Daw et al., 2006). Many tasks also include psychosocial processes, in which participants make altruistic decisions about how to spend money (Hare et al., 2010), or choose to reward/punish themselves over others (Kramer et al., 2007). Other tasks provoke preferences about culturally embedded food brands or luxury items, craft pseudo-stock markets in which participants can make investment decisions and respond to the consequences (Kuhnen & Knutson, 2005), or require participants to move through virtual spaces to collect rewards (Marsh et al., 2010) or escape punishments (Mobbs et al., 2007). Overall, nearly all contemporary reward processing tasks in neuroimaging experiments engage multiple mental operations (Kahnt, 2018), such as working memory (Pochon et al., 2002), rule switching (Remijnse et al., 2009), probability estimation (Hampton, Bossaerts & O’Doherty, 2006), spatial navigation (Marsh et al., 2010), episodic memory (de Greck et al., 2008), emotion regulation (Delgado, Gillis & Phelps, 2008), and/or inhibition (Mengotti, Foroni & Rumiati, 2019).

These research paradigms model a broader array of real-world reward contingencies and have demonstrated that aspects of reward contexts, such as social factors, temporal factors, riskiness, cost or effort required, and the probabilistic nature of contingencies all influence choices and behaviors (Rangel, 2008; Rushworth & Behrens, 2008; Seaman et al., 2018). Internal states, memories of past experiences, and subjective preferences also influence subjective value and choice decisions (Engelmann & Tamir, 2009; Koeneke et al., 2008; Lopez-Persem, Domenech & Pessiglione, 2016). Accordingly, neuroimaging reward processing studies have not only demonstrated RPE-related striatal and mid-brain responsivity (Cooper et al., 2009; Garrison, Erdeniz & Done, 2013; Knutson et al., 2008), but also implicate a wider network of brain activity linked with processing these various peripheral influences (Acikalin, Gorgolewski & Poldrack, 2017; Behrens et al., 2007; Glascher et al., 2010; Juechems et al., 2019; Seaman et al., 2018; Sugrue, Corrado & Newsome, 2004). As such, neurocognitive models of reward learning should endeavor to more fully characterize these additional contextual influences.

Given that reward processing studies utilize a variety of complex tasks, task modeling approaches, and theoretical frameworks, it has been difficult to synthesize a quantitative summary of consistent associations between cognitive constructs and brain function. Emerging meta-analytic techniques have proven helpful in synthesizing neuroimaging findings across disperse experimental sites and diverse scientific approaches (Bottenhorn et al., 2019; Bzdok et al., 2015; Bzdok et al., 2013a; Bzdok et al., 2013b; Clos et al., 2013; Eickhoff et al., 2016; Laird et al., 2015; Ray et al., 2015; Riedel et al., 2018). Utilizing these techniques, we aimed to elucidate brain regions, and in turn networks, consistently linked with different aspects of reward processing.

We employed a data-driven, meta-analytic clustering approach to an extensive body of reward processing neuroimaging results archived in the BrainMap database (www.brainmap.org; (Fox & Lancaster, 2002; Laird, Lancaster & Fox, 2005) to characterize meta-analytic groupings (MAGs) of reward processing experiments based on the spatial similarity of brain activity patterns. These groupings were formed under the assumption that spatially similar task-based brain activity patterns can be categorized as functionally similar, while spatially different patterns can be classified as distinct. To provide insight into the mental processes associated with each of these MAGs, we then conducted functional decoding, a data-driven method for inferring mental processes from observed patterns of brain activity.

While previous reward-related meta-analyses have focused on specific aspects of reward processing (e.g., subjective valuation) (Bartra, McGuire & Kable, 2013; Diekhof et al., 2012; Sescousse et al., 2013; van der Laan et al., 2011), the corpus of neuroimaging results included in our meta-analysis adds to prior work by including brain activity coordinates from a diverse range of tasks and experimental contrasts. The relatively liberal inclusion criteria for our corpus provides a more complete and agnostic investigation of the reward processing construct that arguably better captures its multifaceted nature. Further, while existing reward processing meta-analyses manually grouped published neuroimaging results according to reward-related cognitive constructs of interest to characterize associated brain activity patterns (e.g., Liu, 2011), our meta-analytic, data-driven parsing of experimental contrasts allowed for distinct patterns of brain activity to emerge organically and unbiased by prior assumptions. This *k*-means clustering approach differs from existing meta-analytic strategies as it can identify dissociable patterns of brain activity present within subgroups of studies within the corpus that might otherwise be obscured when focusing only on activity convergence across the entire corpus.

Overall our goals were to: (**1**) provide a data-driven, functionally relevant summary of brain activity patterns commonly observed across diverse reward-related neuroimaging paradigms; (**2**) perform functional decoding analyses linking these activity patterns with distinct cognitive-behavioral processes; and finally, (**3**) to integrate summative interpretations of empirically derived meta-analytic results within existing reward processing theories.

## METHODS

To organize reward-related neuroimaging results into subgroups based on the spatial profile of their modeled activity maps, we performed *k*-means clustering across multiple model orders. We identified viable model orders operationalized as those which maximized between-group differences and minimized within-group differences of between-experiment correlation distributions using four information-theoretic criterion metrics. After comparing and contrasting viable solutions, we ultimately selected the solution with the greatest number of neurocognitively plausible MAGs. Subsequently, convergent brain activity within each grouping of experiments in this selected solution was quantified via separate neuroimaging meta-analyses. Functional decoding assessments using the NeuroSynth database (Yarkoni et al., 2011) (Neurosynth.org) were then performed on the resulting MAG’s unthresholded ALE maps. The following sub-sections elaborate the methodological details of these steps in our meta-analytic clustering approach.

### Identification of included studies

To compile a large corpus of neuroimaging results across reward processing paradigms, we extracted activation coordinates reported in published studies archived in the BrainMap Database as of April 22, 2016, under the paradigm class metadata labels *Reward, Delay Discounting,* and *Gambling* (www.brainmap.org) (Fox et al., 2005; Fox & Lancaster, 2002; Laird et al., 2011). The vast majority (94.9%) of the identified studies were archived under the *Reward* label with most *Delay Discounting* and *Gambling* studies being additionally archived under *Reward*. The *Reward* label denotes that the reported activation coordinates were identified in a task where a stimulus served to reinforce a desired response (e.g., monetary reward after a correct response; www.brainmap.org/taxonomy/paradigms).

We considered only activation coordinates from published neuroimaging studies, among healthy participants, that were reported in standard Talairach (Talairach & Tournoux, 1988) or Montreal Neurological Institute (MNI) (Collins, 1994) space and derived from whole-brain statistical tests. Brain coordinates derived through behavioral correlations or *a priori* region of interest (ROI) analyses were excluded. As this meta-analysis aimed to investigate brain activity linked with typical reward processing, coordinates from groups of individuals diagnosed with neuropsychiatric disorders (e.g., addictive disorders) were excluded from the corpus. Each study included provided at least one experimental contrast that statistically identified brain activity associated with a certain task-event defined by the original authors (e.g., a brain activity map). These experimental contrasts were summarized and curated in the BrainMap database as a set of brain activity foci/coordinates linked either with phases of the original task (i.e., task response, anticipation of outcome, outcome delivery) or stimuli presented in the task (e.g., positive outcome, negative outcome, high reward, low reward). Foci from experimental contrasts could also reflect brain activity locations linked with more abstract and computationally derived constructs of interest in the original study (e.g., learning rate, subjective value).

### Modeled Activation (MA) images of experimental contrasts

Reward-related activation coordinates reported in Talairach space were first transformed into MNI space (Lancaster et al., 2007). Subsequently, modeled activation (MA) images for each experimental contrast (2mm^3^ resolution) were generated by modeling coordinate foci as Gaussian probability distributions to account for spatial uncertainty due to template and between-subject variance (Eickhoff et al., 2009) **Fig. 1, step 1**). This algorithm eliminates the effect of within-experiment number of foci and within-experiment foci proximity by determining the voxel-wise MA values via the maximum probability associated with any one location reported by the experiment instead of taking the union of probabilities associated with all the foci reported by an experiment (Turkeltaub et al., 2012). This ensured that multiple foci from a single experiment did not jointly influence the MA value of a single voxel. Each experimental contrast’s MA image was collapsed to a single dimension vector composed of *v* voxels and then each experiment’s (*e*) vector was concatenated to create an *e* x *v* matrix. The Pearson’s product-moment correlation (*r*) was then computed for each pair of MA maps, resulting in a symmetric *e* x *e* cross-correlation matrix reflecting the extent of similarity (correlation) between the activity locations of different experiments while considering the experiments sample size (**Fig. 1, step 2**).

**Figure 1.**
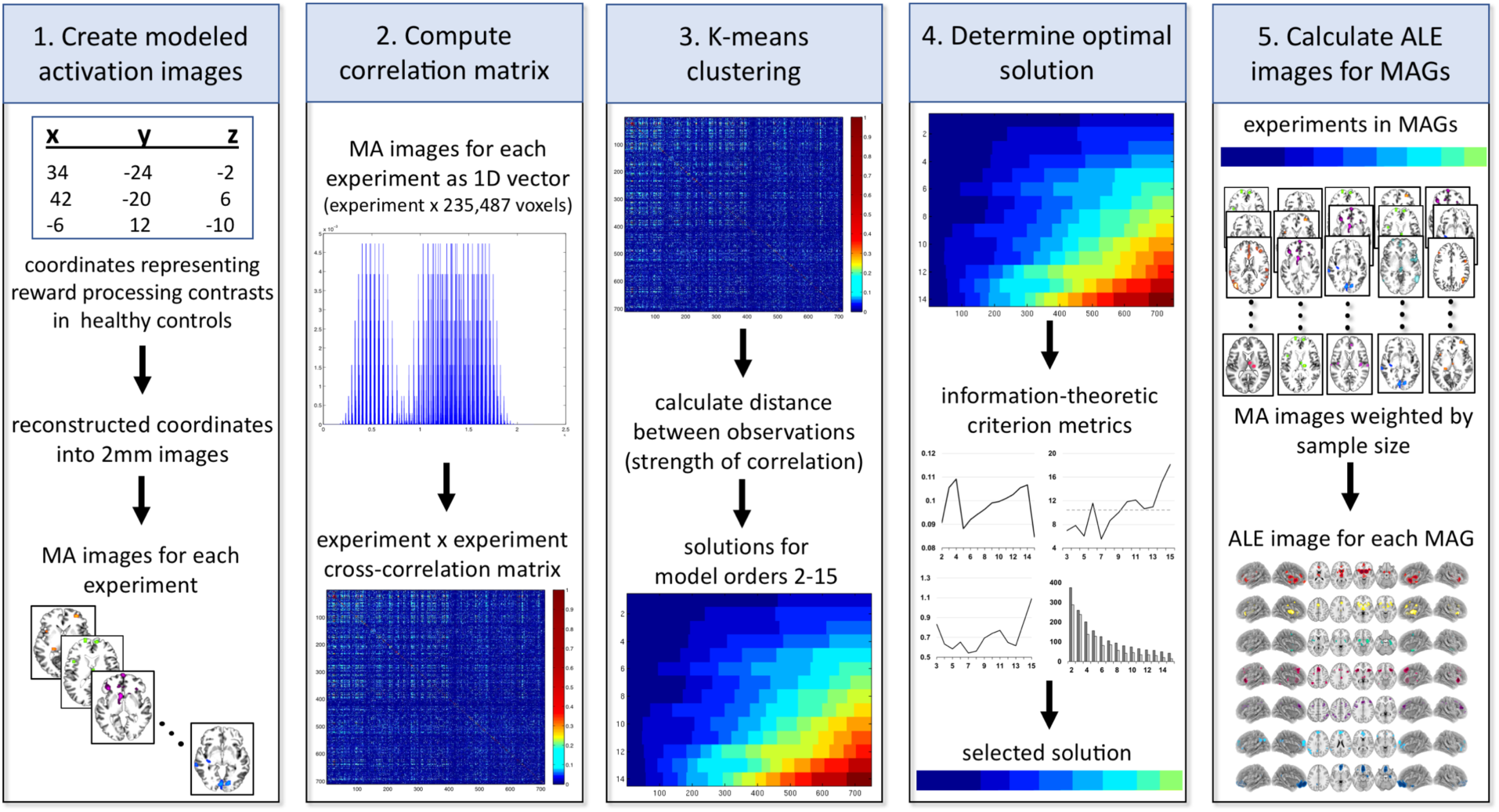
Data extraction and *k*-means clustering analytic workflow. Activation coordinates from experimental contrasts meeting inclusion criteria were extracted from the BrainMap database and reconstructed into modeled activation (MA) images (**step 1**). Each experimental contrast’s MA image was then collapsed to a one-dimensional vector composed of *v* voxels and then each experiment’s (*e*) vector was concatenated to create an *e* x *v* matrix. Pearson’s correlations between every pair of experiment vectors generated an experiment (*e*) x experiment (*e*) cross-correlation matrix (**step 2**). *K*-means clustering analysis grouped experiments according to 14 different model orders (*k* = 2-15; **step 3**). Four different metrics were used to determine viable clustering solutions (**step 4**). Finally, an activation likelihood estimation (ALE) meta-analysis (*p_cluster-level_*<0.05; *p_voxel-level_*<0.001) was performed for each resulting meta-analytic grouping (MAG) of experimental contrasts to compute ALE images of significant activity convergence (**step 5**).

### Correlation matrix-based *k*-means clustering

To group experimental contrasts with similar brain activity patterns, we applied an adapted meta-analytic *k*-means clustering procedure previously applied to large collections of experimental contrasts (Bottenhorn et al., 2019) (**Fig. 1, step 3**). The distribution of experiment-to-experiment distance values, defined as the additive inverse of the Pearson’s correlation coefficient (i.e., *1-r*), was the spatial measure used to determine an experimental contrast’s group assignment. We applied an implementation of *k-*means clustering in the MATLAB environment (version 2014b; Mathworks, Inc.) and classified experiments into *k* groupings. We generated 14 clustering solutions (*k* = 2 through *k* = 15) similar to previous work utilizing *k*-means clustering tehniques on meta-anayltic functional neuroimaging data (Eickhoff et al., 2012; Zangemeister, Grabenhorst & Schultz, 2016). *K*-means clustering is a non-hierarchical clustering approach that segregates observations into a pre-specified number of groups (*k*) (Forgy, 1965; Hartigan & Wong, 1979; Nanetti et al., 2009). This is accomplished by randomly selecting a starting centroid for each group and iterativitly reassigning observations to groups such that within-group variance is minimized and between-group variance is maximized. The random selection of an initial grouping centroid and the subsequent reassignment of experiments was repeated 1,000 times to identify viable solutions (Nanetti et al., 2009).

### Selection of an optimal number of clusters (*k*)

We identified viable model orders based on four information-theoretic criterion metrics that characterized the group separation and group stability properties of each model order (Bzdok et al., 2015; Eickhoff et al., 2016) (**Fig. 1, step 4**). These metrics have previously been used in co-activation-based parcellation analyses to determine optimal clustering solutions (Bzdok et al., 2013b; Chang et al., 2013) for brain regions across task-independent (Bottenhorn et al., 2019; Kahnt et al., 2012; Kelly et al., 2010) and task-dependent (Bzdok et al., 2015; Chase et al., 2015a; Clos et al., 2013; Ray et al., 2015) brain states. We compared and contrasted the outcomes from viable clustering solutions and ultimately selected the solution identifying the greatest number of neurocognitively plausible MAGs for presentation in the main text. The outcomes of other viable clustering solutions are presented in the supplemental materials.

Regarding between-group separation, solutions indicating greater separability were desired as they possess a lower likelihood of experimental contrast misclassification into neighboring groupings. Two metrics were used to assess group separation. The first was the average silhouette coefficient (ASC) (Kaufman & Rousseeuw, 2009) which quantified a given experiment’s similarity to the others within its designated grouping relative to the experiments in other groupings. This criterion metric identifies viable solutions (e.g., *k*) as those with significantly higher coefficients than the subsequent solution (*k*+1) or as solutions without significantly lower coefficients than the preceding solution (*k*-1). The second separability metric was the hierarchy index (HI) which compared the percentage of experiments assigned to a given grouping that were lost from that grouping when moving from the *k* to the *k*-1 solution for all solutions greater than *k* = 2 (Kahnt et al., 2012). Experiments are generally thought to be lost from a grouping due to incorrect group assignment. Viable solution candidates were defined as those with a percentage of lost experiments below the median value for all possible solutions.

Regarding within-group stability, solutions demonstrating higher stability were desired as high stability indicates consist and robust grouping. Two metrics were employed to assess group stability. The first was the variation of information (VI) statistic (Meilă, 2007) which described the amount of information lost or gained when moving to a subsequent clustering solution. Viable solutions were defined as those where the VI increased from the *k* to *k*+1 solution (primary criterion) and where the VI decreased from the immediately preceding *k*-1 solution to the *k* solution (secondary criterion) (Clos et al., 2013). The second metric used to assess group stability was the minimum number of experimental contrasts consistently assigned to a grouping, relative to the mean number of experimental contrasts consistently assigned to that grouping across the 1000 iterations (Nickl-Jockschat et al., 2015; Poldrack, 2006). Viable solution candidates were defined as those with a minimum number of consistently assigned experiments that was greater than half the mean number of consistently assigned experiments.

### Statistical convergence within meta-analytic groupings

The *k*-means clustering algorithm assigns all the experimental contrasts within the corpus to separate collections/groupings. We refer to the collections of experimental contrasts in the *k*-means clustering solutions as meta-analytic groupings (MAGs). After identifying a clustering solution, convergent brain activity maps across all experimental contrasts within each of the *k* MAGs were produced using a revised activation likelihood estimation (ALE) algorithm implemented in the MATLAB environment (**Fig. 1, step 5**) (Eickhoff et al., 2012; Eickhoff et al., 2009; Turkeltaub et al., 2012). Voxel-wise ALE scores were computed as the union of all MA images within each MAG and served as a quantitative representation of convergent activity within a grouping. Each MAG’s distribution of ALE scores was then subjected to significance testing utilizing a null distribution analytically derived from the random spatial association between experiments (Eickhoff et al., 2012). The resulting *p*-values were thresholded at *p_cluster-level_*<0.05 (FWE corrected for multiple comparisons; cluster-forming threshold *p_voxel-level_*<0.001) to identify significant clusters within each MAG. Convergent activity maps were exported to MANGO (http://ric.uthscsa.edu/mango/) and rendered for visualization utilizing Nilearn 0.5.0 (Abraham et al., 2014).

### Functional decoding of meta-analytic groupings

To gain insight into cognitive and behavioral processes associated with convergent brain activity captured within each MAG, we performed functional decoding utilizing NeuroVault (Gorgolewski et al., 2015) and NeuroSynth (Yarkoni et al., 2011). NeuroVault is a web-based repository allowing researchers to store, share, and visualize statistical neuroimaging maps. NeuroVault also facilitates functional decoding by interfacing with NeuroSynth, a platform that produces term-specific meta-analytic maps based on its database containing activation coordinates and automatically extracted terms from over 14,000 published neuroimaging studies (abstracts). Functional decoding was then performed by uploading each MAG’s unthresholded ALE map to NeuroVault which then correlated these uploaded maps with term-specific meta-analytic maps extracted from NeuroSynth’s database. This yielded a ranked list of maximally related NeuroSynth (NS) terms and their respective Pearson correlation values. These correlation values represented how well the spatial distribution of activity associated with each term in the database matched the activity pattern of the uploaded MAG’s ALE map. This functional decoding approach allowed for an objective description of the NS terms linked with each MAG, as well as a comparison of our MAGs with the broader neuroimaging literature.

## RESULTS

The search criteria yielded a corpus of brain activation foci from a total of 749 experimental contrasts across 176 studies involving 13,358 healthy participants. Published articles with experimental contrasts included in the corpus are listed in **Table S1**. Of these 749 contrasts, 711 (94.9%) were archived under BrainMap’s *Reward* paradigm class label, 42 (5.6%) under the *Delay Discounting* paradigm class label, and 20 (2.7%) under the *Gambling* paradigm class label. Almost all studies included in the corpus were also archived under a variety of other meta-data labels (e.g., *Task Switching* [6.4%], *Go/No-Go* [2.9%], *Visuospatial Attention* [2.9%], *Wisconsin Card Sorting Test* [2.6%], *Reasoning/Problem Solving* [1.3%]), which is unsurprising as reward processing is a multifaceted construct, connecting elements of sensation, perception, cognitive control, and other mental operations. As studies included in the corpus could be dually archived under multiple paradigm classes, the percentages of corpus experiments in each paradigm class was not expected to sum to 100%. A complete list of the percentage of experiments achieved under each paradigm class is presented in **Table S2**.

The foci included in our corpus were from 749 experimental contrasts that probed specific events from a variety of tasks including, but not limited to: monetary incentive delay (MID) tasks; gambling simulation tasks (e.g., wheel of fortune, four-arm bandit/slot machine, blackjack tasks); stock market simulation tasks; delay discounting tasks; face, product, or food preference tasks; punishment avoidance tasks (i.e., predator, prey paradigm); reversal learning tasks (e.g., Wisconsin card sort tasks); Iowa gambling tasks; tower of London tasks; go/no-go tasks; and balloon analogue risk tasks (BART). However, not all studies utilized identical versions of these well-known paradigms and even studies applying the same task often reported brain activity from different experimental contrasts.

### Clustering Solutions

To identify viable clustering solutions, we quantitatively evaluated solutions *k* = 2 through *k* = 15 across four metrics: the average silhouette coefficient (ASC), the hierarchy index (HI), the variation of information (VI) statistic, and a cluster-size metric (**Fig. 2**). The ASC metric identified the *k* = 4 model order as a viable solution as it had a higher ASC than the subsequent *k* = 5 model order, but not a lower ASC than the preceding *k* = 3 model order (**Fig. 2A**). Regarding HI, model orders *k* = 3, 4, 5, 7, 8, and 9 met criterion, having a percentage of lost experiments below the median with the *k* = 7 model order having the lowest percentage of lost experiments (**Fig. 2B**). Regarding VI, the *k* = 5, 7, and 13 model orders all met primary and secondary metric criteria. Specifically, the subsequent solution and the immediately preceding solution both displayed an increased VI statistic (**Fig. 2C**). Regarding the cluster consistency metric, all solutions except *k* = 14 and *k* = 15 had a minimum number of consistently assigned experiments that was greater than half the mean number of consistently assigned experiments indicating high cluster consistency (**Fig. 2D**).

**Figure 2.**
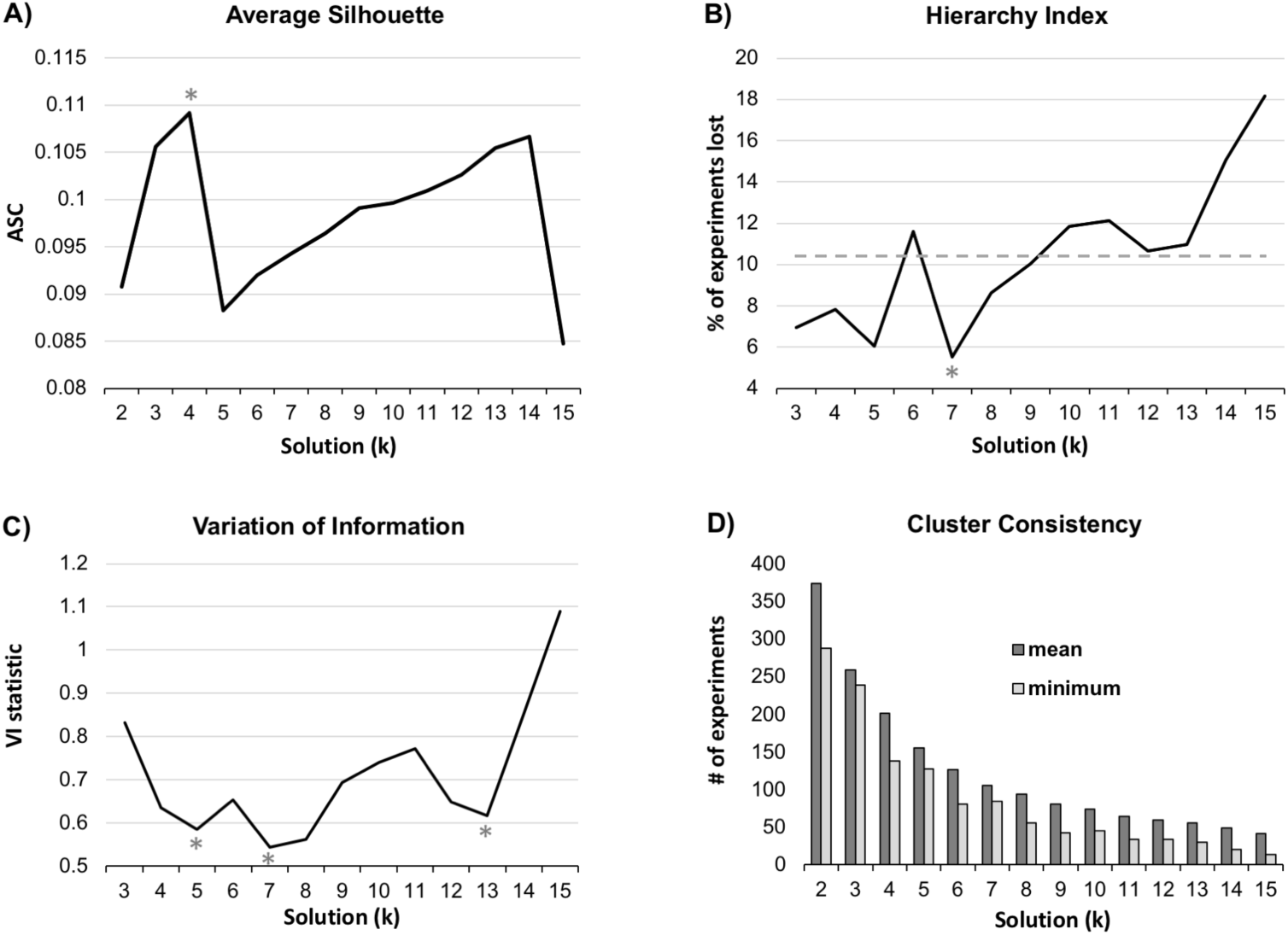
Clustering solution metrics for *k* = 2 - 15. (**A**) A higher average silhouette coefficient (ASC) indicates increased group-separation and a lower likelihood of experiment misclassification. The ASC identified the *k* = 4 model order as a viable solution. (**B**) However, the hierarchy index (HI) indicated that the *k* = 7 model order not only displayed a below-median (gray dashed line) number of experiments lost from the previous model order, but displayed the lowest HI relative to all other model orders. (**C**) The *k* = 5, 7, and 13 model orders met both the primary and secondary criterion for the variation of information (VI) metric as they all displayed an increased VI statistic moving from *k* to *k*+1 and displayed a decreased VI statistic moving from *k*-1 to *k.* This indicated that the *k* = 5, 7, and 13 model orders had high group-stability. (**D**) All solutions except *k* = 14 and *k* = 15 had a minimum number of consistently assigned experiments that was greater than half the mean number of consistently assigned experiments indicating high cluster consistency. Given that model orders *k* = 4, 5, and 7 were identified across multiple metrics, we considered these to be viable clustering solutions. (* indicates the best solution(s) for each metric).

Based on these metrics we identified model orders *k* = 4, 5, and 7 as viable solutions as they all met criteria for three of the four metrics. Ultimately, we selected the *k* = 7 solution for presentation in the main text as it identified the greatest number of neurocognitively plausible MAGs. However, we also performed supplementary analyses and examination of the *k* = 4 and *k* = 5 solutions. Clustering (**Fig. S1 & S3**) and functional decoding results (**Fig. S2 & S4, Table S4-5**) for these two other solutions are presented in the supplemental material and discussed in relation to the *k* = 7 outcomes. We suggest that examination and functional decoding of these three viable model orders provides additional information regarding the integration and segregation of functional brain networks and cognitive-behavioral constructs across varying levels of parcellation.

### Meta-Analytic Groupings

As depicted in the experiment (*e*) x experiment (*e*) cross-correlation matrix ordered by the seven groupings (**Fig. S5**), MAGs from the *k* = 7 clustering solution displayed between-MAG differences and within-MAG similarities when considering pairwise experiment-experiment correlation distributions. Further, each of the 7 MAGs contained a reasonably high number of experiments and foci from studies involving a large pool of participants (**Table 1**). The generated ALE maps identified regions of convergent activity for each of the seven MAGs (*p_cluster-level_* < 0.05; FWE-corrected; *p_voxel-level_* < 0.001; **Fig. 3, Table S3**). The order in which the seven derived MAGs were labeled does not reflect any aspect of the *k*-means clustering analysis.

**Figure 3.**
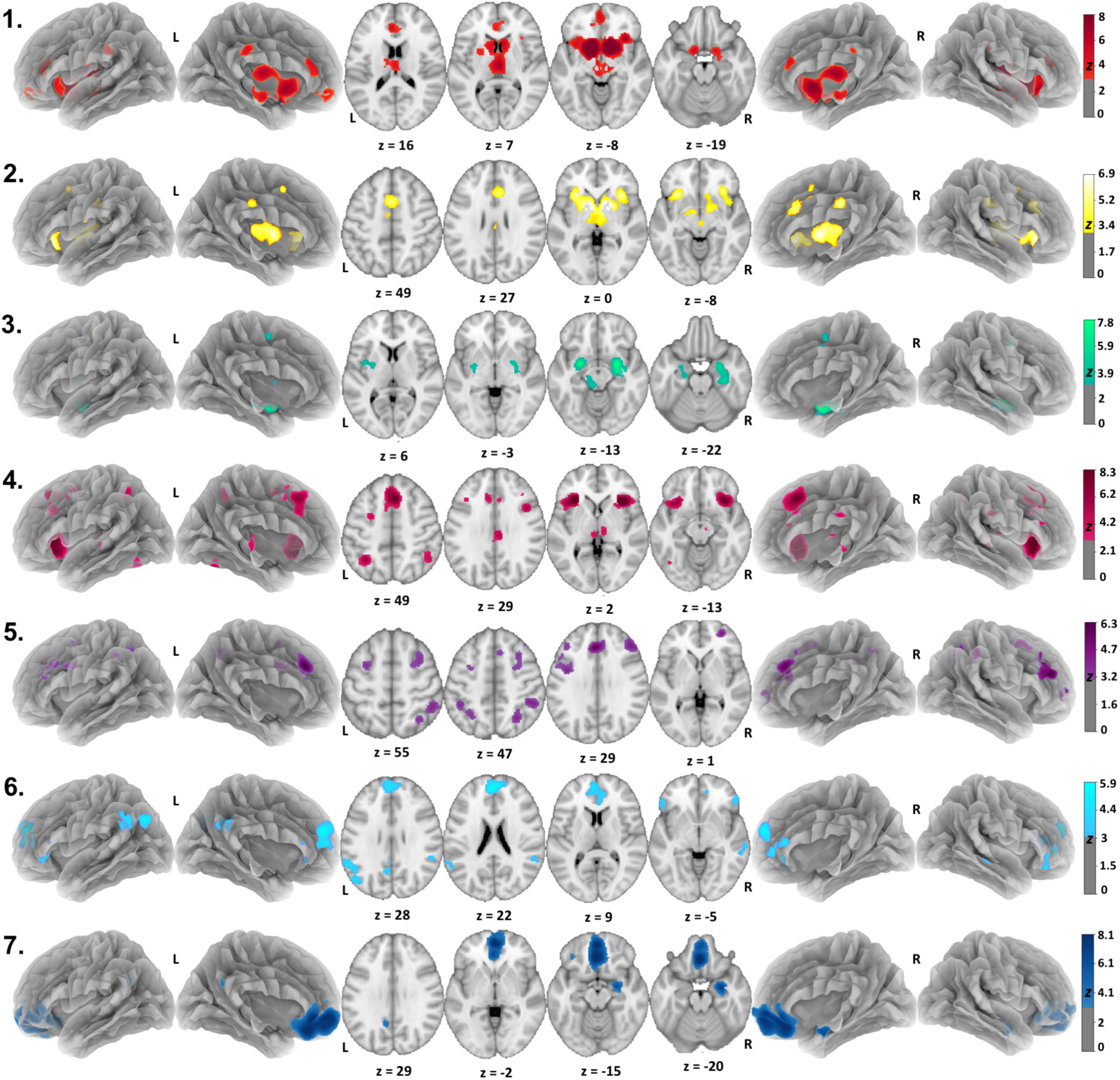
Brain activity profiles associated with each meta-analytic grouping (MAG) of reward processing experiments derived via *k*-means clustering. ALE images identified significant (*p_cluster-corrected_* < 0.05; *p_voxel-level_*< 0.001) convergence in dissociable and distributed brain regions across each MAG. Unthresholded maps of each MAG are available on NeuroVault (https://neurovault.org/collections/5070/).

**Table 1.** Number of experiments, subjects, and foci within each MAG from the *k* = 7 clustering solution.

| MAG | Experiments | Subjects | Foci |
| --- | --- | --- | --- |
| MAG 1 | 181 | 3,226 | 1,609 |
| MAG 2 | 99 | 1,583 | 1,101 |
| MAG 3 | 89 | 1,717 | 601 |
| MAG 4 | 98 | 1,775 | 886 |
| MAG 5 | 94 | 1,766 | 667 |
| MAG 6 | 88 | 1,652 | 546 |
| MAG 7 | 100 | 1,639 | 541 |
| **Total** | **749** | **13,358** | **5,951** |

The ALE map for <u>MAG-1</u> (**Fig. 3, red**) displayed bilateral convergent activity in a large cluster extending through most of the striatum and thalamus and into the mid-brain and anterior insula with the center of mass in the ventral striatum (VS). MAG-1 also included smaller clusters in the subgenual anterior cingulate, posterior cingulate cortex (PCC), and ventromedial prefrontal cortex (vmPFC). <u>MAG-2</u> (**Fig. 3, yellow**) also exhibited convergent activity in the striatum, thalamus, medial prefrontal cortex (mPFC), cingulate, and anterior insula. Whereas MAG-1 included activity in the VS, vmPFC, anterior cingulate cortex (ACC) and PCC, MAG-2 displayed convergent activity centered on more dorsal regions (**Fig. S6A**), specifically in the dorsal striatum (DS), dorsal medial prefrontal cortex (dmPFC), pre-supplementary motor area (pre-SMA), and dorsal medial cingulate cortex (dmCC). <u>MAG-3</u> (**Fig. 3, green**) exhibited bilateral convergent activity in the amygdala and hippocampus, as well as in the SMA and left posterior insula. <u>MAG-4</u> (**Fig. 3, pink**) displayed convergence in the bilateral insulae and the dmPFC extending into the dorsal ACC. MAG-4 also included smaller bilateral clusters in the posterior thalamus encompassing the epithalamic habenular nuclei and in the lateral PFC (LPFC), PCC, and the bilateral intraparietal sulcus (IPS). <u>MAG-5</u> (**Fig. 3, purple**) also displayed convergence in the bilateral IPS extending into the inferior and superior parietal lobules, dmPFC, and the bilateral LPFC including clusters in the dorsal middle frontal gyrus and frontal pole. <u>MAG-6</u> (**Fig. 3, aqua**) displayed convergence in the central medial PFC (cmPFC), subgenual ACC, right middle temporal gyrus, bilateral inferior frontal gyrus, precuneus, and bilateral temporoparietal junction (TPJ) extending into the right middle temporal gyrus and left supramarginal gyrus. <u>MAG-7</u> (**Fig. 3, navy**) also displayed convergence in the medial frontal cortex, but ventral to that for MAG-6 (**Fig. S6B**). Specifically, MAG-7 displayed a large cluster extending across the entire medial orbital frontal cortex (OFC) and subgenual ACC in addition to convergence in the right amygdala, right hippocampus, PCC, and left middle frontal gyrus.

### Functional Decoding

NeuroSynth functional decoding results were exported as term correlation values quantifying the similarity between an input map (i.e., each MAG’s ALE map) and maps associated with the NS term. To facilitate interpretation, the top 10 NS terms with the highest correlation values for each MAG were used to generate behavior profiles (**Fig. 4, Table**

**Figure 4.**
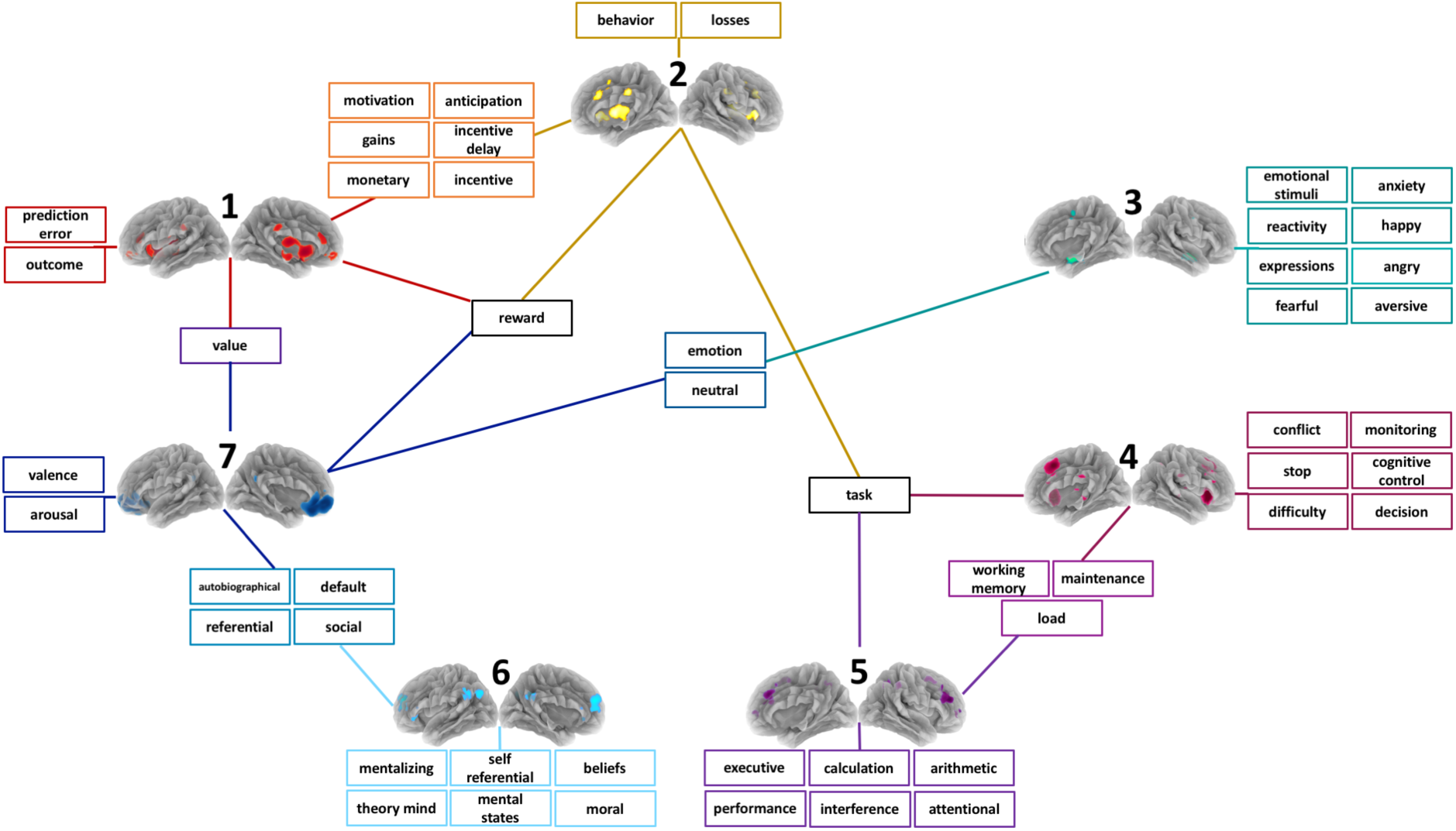
Behavior profiles associated with each meta-analytic grouping (MAG) of reward processing experiments. Behavior profiles consisted of NeuroSynth (NS) terms with the top 10 highest correlation values for each MAG (excluding anatomical terms). Lines connect MAGs to the terms making-up their unique behavior profile. Additionally, some terms or groups of terms are connected to multiple MAGs indicting that these terms were in the top 10 lists of multiple MAG’s.

**3).** Anatomical NS terms, such as “insula” or “anterior,” were excluded from these lists. NS terms that were near duplicates of those already on the list were condensed into the one term that was highest on the list. For example, the terms “emotional” and “emotions” would be condensed if the term “emotion” was already in the top 10 (**Table 3**, Near Duplicates section).

**Table 3.**
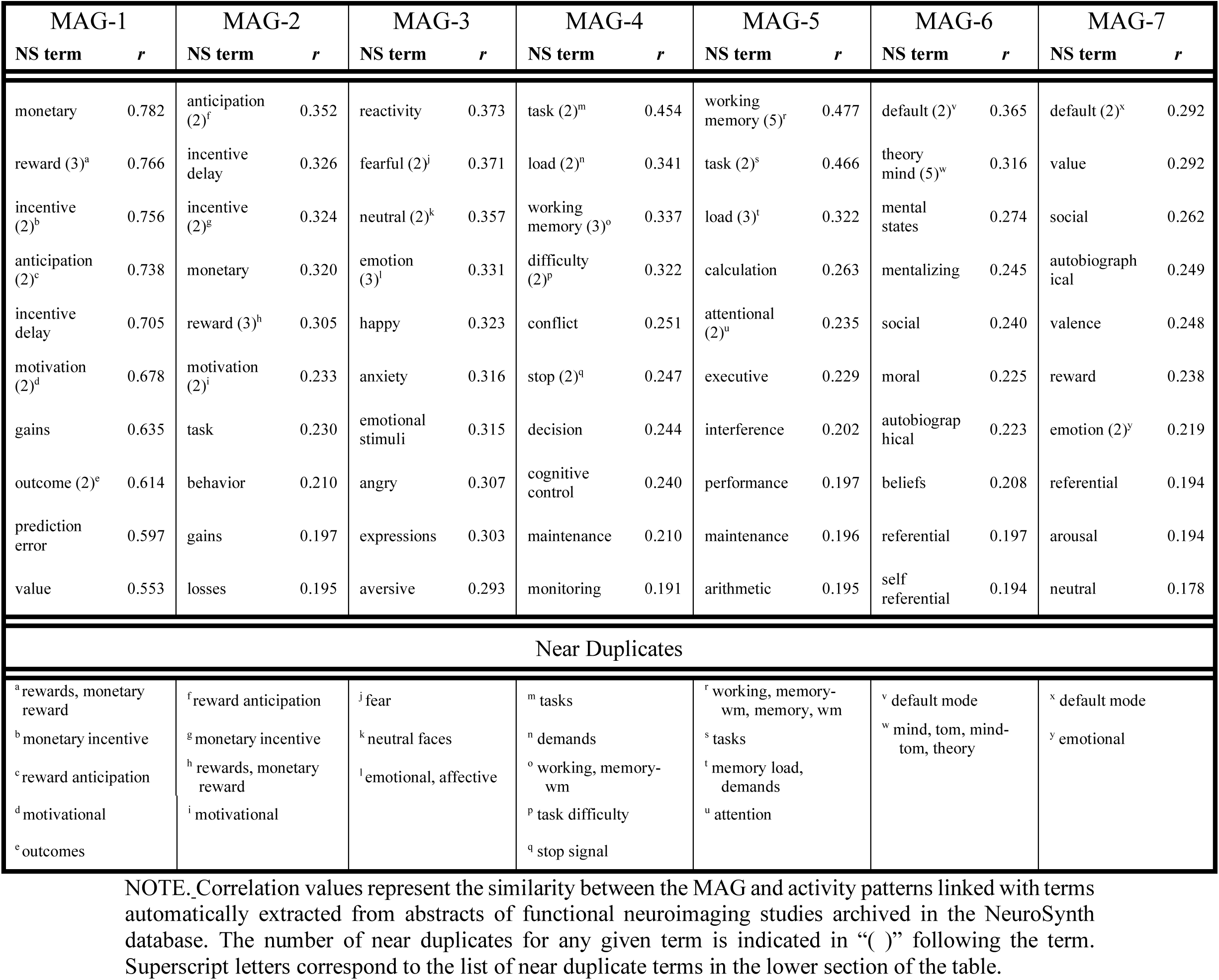
NeuroSynth (NS) terms composing each MAG’s behavior profile and their respective correlation values.

### Functional Interpretations of MAGs

We provide an interpretation of each MAG’s presumed functional role based on formal decoding outcomes and previous studies involving the brain regions displaying convergent activity. These functional interpretations speak to probabilistic co-occurrence of activity patterns with certain cognitive-behavioral functions across the literature; but they do not define the function of specified brain locations, per se.

**MAG-1** displayed convergent activity in the mid-brain, vmPFC, and VS (**Fig. 3, red**). The VS and its dopaminergic connections are known to respond to reward cues and unexpected outcomes or reward prediction errors (RPEs) (Hollerman & Schultz, 1998; O’Doherty et al., 2003; Parker et al., 2016; Schultz, Dayan & Montague, 1997). Functional decoding results indicated that MAG-1 was associated with the terms *prediction error*, *outcome*, *anticipation, reward, gains,* and *incentive* (**Fig. 4, Table 3**). These terms, in conjunction with the convergent activity observed in the VS and vmPFC, suggest that MAG-1 may play a role in processing reward cues, unexpected outcomes, and **encoding RPEs**.

**MAG-2** also showed convergent activity in the striatum and mPFC, albeit in more dorsal regions than those observed for MAG-1. Specifically, MAG-2 displayed activity in the dmPFC/pre-SMA, DS, dmCC, and anterior insula (**Fig. 3, yellow)**. Behaviorally, MAG-2 was associated with many of the same terms as MAG-1 suggesting RPE encoding and reward cue processing, specifically, *anticipation*, *reward*, *gains*, *incentive, incentive delay*, *monetary*, and *motivation* (**Fig. 4, Table 3**). In addition, MAG-2 was also associated with three other terms: *losses*, *behavior,* and *task*. Evidence suggests that more dorsal regions of the striatum, mPFC, cingulate, and the pre-SMA may specialize in processing response-outcome contingencies (Atallah et al., 2007; McNamee et al., 2015; O’Doherty et al., 2004b; Parker et al., 2016) and negative outcomes (Bartra, McGuire & Kable, 2013; Tops & Boksem, 2012). MAG-2’s brain activity profile and behavioral profile were consistent with this view and suggest involvement in RPE encoding, but perhaps with a specialization for **response-outcome contingencies** and/or involving **negative outcomes.**

**MAG-3** exhibited convergent activity in the amygdala, hippocampus, left posterior insula, and SMA (**Fig. 3, green**). Previous research has implicated these brain regions in the recognition and regulation of emotions (Adolphs, Baron-Cohen & Tranel, 2002; Grecucci et al., 2013; Phan et al., 2003; Phan et al., 2004; Schnell et al., 2011; Seymour & Dolan, 2008). Our functional decoding results were consistent with these previous findings as MAG-3’s behavior profile included affective terms such as *reactivity*, *emotion*, *emotional stimuli*, *anxiety*, *fearful*, and *happy* (**Fig. 4, Table 3**). In conjunction with the brain activity profile, one functional interpretation of MAG-3 is a role in **emotional processing**.

**MAG-4** included convergent activity in the bilateral anterior insulae, daCC, dmPFC, and thalamus with clusters encompassing the epithalamic habenular nuclei (**Fig. 3, pink**). The anterior insula, dACC, and thalamus represent core nodes of the salience network (SN) (Seeley et al., 2007), a commonly observed large-scale brain network thought to be involved in detecting salient stimuli and, in turn, adjusting attentional focus by modulating the relative activity level of other networks (Menon & Uddin, 2010; Sridharan, Levitin & Menon, 2008; Sutherland et al., 2012). MAG-4’s behavior profile included terms indicating a role in generalized ‘task-on’ processes (e.g., *cognitive control*, *decision*, *task*), and both MAG-4 and MAG-5 were jointly associated with terms related to the specific executive function working memory (*working memory*, *maintenance*, *load*; **Fig. 4, Table 3**). Further, MAG-4’s behavior profile also included the term *stop* suggestive of another executive function, specifically, inhibitory control/response inhibition. MAG-4 convergent activity also encompassed the habenula, an epithalamic nucleus implicated in conflict monitoring, error processing, reward, and consequently decision-making (Baker et al., 2016; Ide & Li, 2011; Kawai et al., 2015). MAG-4’s pattern of convergent activity resembled previous accounts of human habenula connectivity (Ely et al., 2019; Ely et al., 2016) particularly during error processing (Ide & Li, 2011) and thus, it is noteworthy that MAG-4’s behavior profile also includes terms related to performance monitoring (*conflict*, *monitoring*, *and cognitive control*). In conjunction with the observed convergent activity in the anterior insula, dACC, and habenula, the collection of terms in MAG-4’s behavior profile suggested a role in **externally-focused attention** particularly in the context of **executive functions** such as working memory, response inhibition, and performance monitoring.

**MAG-5** presented convergent activity in the LPFC and parietal cortex (**Fig. 3, purple**), core nodes of the executive control network (ECN) (Seeley et al., 2007). The ECN is thought to play a prominent role in attention, inhibition, cognitive flexibility, working memory, and planning (Brown et al., 2019; Niendam et al., 2012) and has also been linked with reward motivated decision-making (Hobkirk et al., 2019; Tanji & Hoshi, 2008). As noted above, MAG-5 and MAG-4 were jointly associated with terms related to working memory (*working memory*, *maintenance*, *load*; **Fig. 4, Table 3**) and similar to MAG-4, MAG-5’s behavior profile also included terms that indicated a role in generalized ‘task-on’ processes (*executive*, *attentional, performance*). Additionally, MAG-5 was associated with terms indicative of mathematical calculating (*calculation*, *arithmetic*). Overall, this collection of terms suggested a role in **externally-focused attention** particularly in the context of **higher-order mental operations** such as mathematical calculation.

**MAG-6** displayed convergent activity in the cmPFC, parietal lobe, TPJ, middle temporal gyrus, PCC, and precuneus (**Fig. 3, aqua**), regions resembling the default-mode network (DMN). The DMN is a commonly observed collection of brain regions showing decreased activity during overt task performance (Raichle et al., 2001) and increased activity during ‘rest’ (Spreng, 2012; Spreng & Grady, 2009) or internally-focused tasks involving theory of mind, autobiographical memory retrieval, or other forms of self-referential thought (Andrews-Hanna, 2012; Mars et al., 2012; Qin & Northoff, 2011; Spreng & Grady, 2009). MAG-6’s behavioral profile was consistent with these functions previously linked to the DMN and included terms suggesting self-referential thought (*autobiographical*, *self-referential, beliefs*), mentalizing (*beliefs*, *mental states*, *mentalizing*), and theory of mind (*theory mind, social, moral*; **Fig. 4, Table 3**). The convergent activity in DMN regions, in conjunction with this collection of terms suggested involvement of this MAG in **internally-focused thought** specifically regarding **abstract mental states**.

**MAG-7** also displayed convergent activity in DMN regions (**Fig. 3, navy**), including the medial OFC, ACC, and PCC (Acikalin, Gorgolewski & Poldrack, 2017; Raichle et al., 2001; Spreng, 2012; Spreng & Grady, 2009) However, the convergent activity in the medial OFC and PCC was located more ventrally than that from MAG-6. Additionally, MAG-7 did not display convergent activity in the TPJ or temporal gyrus, as in MAG-6, but instead displayed convergence in the right amygdala. Interestingly, the vmPFC, medial OFC, amygdala, and PCC have been implicated in subjective valuation and have been previously characterized as constituting a subjective valuation network (Bartra, McGuire & Kable, 2013; Levy & Glimcher, 2012; Peters & Buchel, 2010). While both MAG-7 and MAG-6 were jointly associated with terms suggesting intrapersonal and interpersonal mentalizing (e.g., *autobiographical*, *social*, *referential*), MAG-7 was also associated with terms related to subjective valuation and arousal (i.e., *value*, *valence*, *arousal*, *reward*, *neutral,* and *emotion*) (**Fig. 4, Table 3**). Overall the localization of convergent activity in conjunction with the functional decoding of MAG-7 suggested a role in **internally-focused thought** specifically in the context of **subjective valuation.**

## DISCUSSION

Using a data-driven *k*-means clustering approach, we classified 749 reward processing experiments into seven meta-analytic groupings (MAGs). We then computed and visualized the convergent brain activity for each MAG in the ALE framework. Our results suggested that the underlying organization of brain activity during reward processing tasks could be dissociated into separate networks each with distinct functional roles. By performing exploratory functional decoding analyses, we constructed a behavior profile for each MAG comprised of cognitive-behavioral terms that captured common and unique mental operations that may be critically linked with each MAG’s brain activity profile. We note that while our functional decoding results suggest differential functioning across brain activity patterns, they do not definitively prescribe functional specializations to these patterns.

### MAG functions within a reward processing framework

To serve as a roadmap for the discussion below, we conceptualized these outcomes in a reward processing heuristic framework. This framework, combining the summative interpretations of all MAGs, captures cognitive mechanisms involved with various potential influences on reward processing present in the complex contexts generated by various reward processing tasks (**Fig. 5**). In the more complex situations generated by human neuroimaging paradigms, we propose that value predictions, based on RPE signals generated through stimuli-outcome and response-outcome pairings, are further influenced by external contextual factors (Behrens et al., 2007; Fiorillo, Tobler & Schultz, 2007; Hariri et al., 2006; Porcelli & Delgado, 2009; Preuschoff & Bossaerts, 2007; Rangel, 2008; Seaman et al., 2018), and/or by subjective internal factors (Hetherington, Pirie & Nabb, 2002; Juechems et al., 2019; Lopez-Persem, Domenech & Pessiglione, 2016; Singer, Critchley & Preuschoff, 2009).

**Figure 5.**
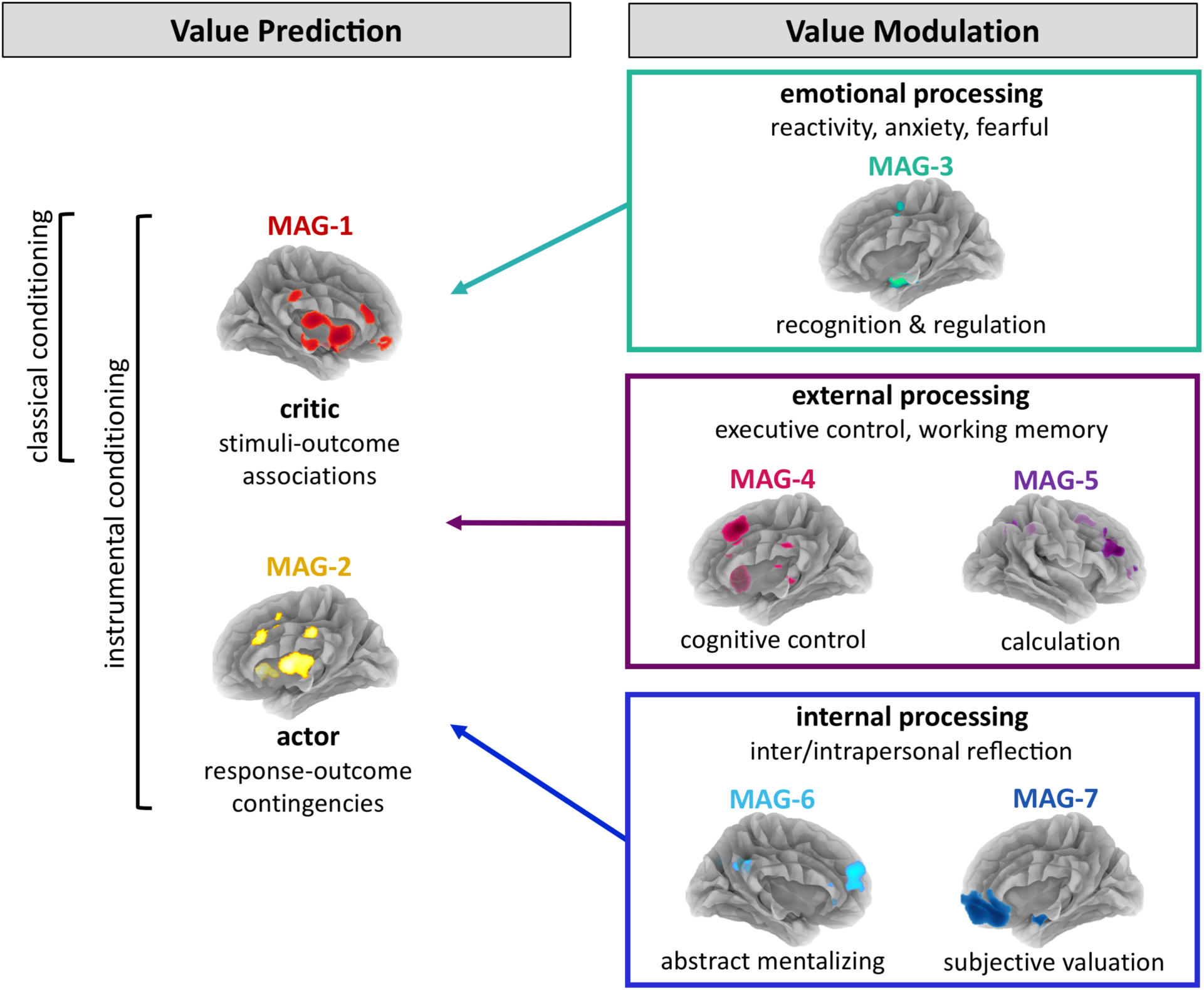
Heuristic framework linking each MAG’s unique functional interpretation. This schematic depicts a synthesized interpretation of each MAG’s dissociable functions based on term definitions and previously implicated cognitive processes. We propose that critic (ventral) and actor (dorsal) striatal-medial frontal pathways form value predictions through classical conditioning (stimuli-outcome associations) and/or instrumental conditioning (response-outcome contingencies). These value predictions are further modulated by a variety of emotional factors (stress, fear, anxiety), external factors (task calculations, task performance, risk, delay, effort, probability), and other internal factors (subjective preferences, personal beliefs, internal states, memories of past experiences). The integration of these influences results in a final representation of value that ultimately governs behavior.

While MAG-1 and MAG-2 identified convergent brain activity in a network of regions implicated in value prediction through RPE signals during classical and instrumental conditioning (an interpretation further supported by quantitatively defined behavior profiles), the other five MAGs possessed behavior profiles suggesting putative roles in processing more peripheral, external, and internal influences present in the complex reward contingencies inherent to more externally valid reward learning circumstances. Based on these results, our heuristic framework proposes that emotional, external, and internal influences in a given context play a role in modulating the valuation of outcomes formed and updated by RPE signals (**Fig. 5**). The following subsections describe each MAG’s interpretation in more detail within this conceptualization.

### Value prediction through RPE encoding

The most basic elements of value prediction are often studied within conditioning paradigms in which a reward or punishment is paired with a neutral stimulus (O’Doherty, 2004a; Porcelli & Delgado, 2009). After this association has been learned, the neutral stimulus acquires a predictive value (Hollerman & Schultz, 1998; Schultz, Dayan & Montague, 1997). Neuroimaging findings have provided robust evidence of dopaminergic signals in the striatum encoding RPEs (O’Doherty, 2004a; O’Doherty et al., 2003; Porcelli & Delgado, 2009; Wang, Smith & Delgado, 2016). These signals are thought to form value predictions and learned stimulus-outcome and response-outcome associations (Delgado et al., 2000; Gottfried, O’Doherty & Dolan, 2003; Hollerman & Schultz, 1998; Knutson et al., 2001; Schultz, 2016; Zald et al., 2004). Additionally, blood oxygen-level dependent (BOLD) activity in the mPFC, thalamus, mid-brain, amygdala, insula, and inferior frontal cortex has been associated with RPE-related task events (Chase et al., 2015b; Fouragnan, Retzler & Philiastides, 2018; Garrison, Erdeniz & Done, 2013; Rutledge, 2010). Both of our MAGs that included striatal activity (i.e., MAG-1, MAG-2; in conjunction with other regions) exhibited behavior profiles consistent with RPE task event processing.

That said, accumulating evidence suggests that there may be dissociable ventral and dorsal RPE-encoding pathways with distinct functional roles (García-García, 2017; O’Doherty et al., 2004b; Parker et al., 2016) and that these roles can be mapped to a well-studied computational framework termed the actor-critic model (García-García, 2017; Niv, 2009). Specifically, the pathway from the ventral tegmental area (VTA) to the VS to limbic and cortical regions (i.e., the mesocorticolimbic pathway) is thought to use RPEs to update stimuli values, corresponding to the role of the critic in actor-critic models. Whereas the pathway from the substantia nigra (SN) to the DS to the motor and premotor cortex (i.e., the nigrostriatal pathway) is thought to assign this value to actions and be involved in action-selection, corresponding to the actor role in actor-critic models (Barto, 1995; García-García, 2017).

The distinction between regions displaying convergent activity in MAG-1 and MAG-2 is in line with such actor-critic models distinguishing between ventral and dorsal pathways. MAG-1 displayed convergent activity in the striatum, thalamus, mid-brain, amygdala, anterior insula, and ventral aspects of the mPFC and the associated behavioral profile included the terms *prediction error*, *anticipation*, *reward, outcome,* and *incentive* supporting previous accounts of these regions responding to reward cues and RPEs (Bray & O’Doherty, 2007; Kirsch et al., 2003; McClure, Berns & Montague, 2003). MAG-2, while also displaying convergent activity in the striatum and mPFC, was localized to more dorsal regions and included activity in the dmCC and pre-SMA (**Fig. S6A**). MAG-2 was also associated with terms indicative of RPE encoding, specifically *incentive*, *reward*, *losses*, *motivation*, and *anticipation*. However, unlike MAG-1’s behavioral profile, that for MAG-2 included the terms *behavior* and *task* suggestive of learning more instrumental, response-outcome contingencies. Evidence suggests that DS function is also linked to RPE encoding and particularly so during instrumental conditioning paradigms (Atallah et al., 2007; Elliott, 2004; García-García, 2017; Hart, Leung & Balleine, 2014; Haruno, 2004; O’Doherty et al., 2004b). In these paradigms, outcomes are contingent on the execution of an action. As instrumental contingencies are learned, actions that lead to desired outcomes are more likely to be repeated. Neuroimaging studies investigating this instrumental component of conditioning support the notion that DS activity may reflect dopaminergic signaling underlying response-outcome contingency learning similar to the way the ventral striatum supports stimulus-outcome learning (Atallah et al., 2007; Haruno, 2004; O’Doherty et al., 2004b). Further, the dmCC and pre-SMA have been shown to regulate performance of instrumental, goal-directed responses (Liljeholm, Dunne & O’Doherty, 2015; Mannella, Mirolli & Baldassarre, 2016; McNamee et al., 2015). Thus, the convergent activity observed in the dmCC, pre-SMA, and DS in MAG-2 corresponds well with regions previously implicated in instrumental conditioning. In addition to positive outcomes, this network is also thought to specialize in processing negative outcomes (Bartra, McGuire & Kable, 2013). Overall, our data-driven, meta-analytic groupings support the contemporary notion of functionally dissociable ventral and dorsal RPE encoding networks. Specifically, MAG-1 captured a network including VS and vmPFC activity that previous research has implicated in representing and updating value in stimuli-outcome associations, while MAG-2 captured a network including the DS, dmCC, pre-SMA, and the anterior insula implicated in processing response-outcome contingencies and negative outcomes (Bartra, McGuire & Kable, 2013; O’Doherty et al., 2004b).

### Value modulation by dissociable influences

The behavior profiles associated with MAG-3-7 suggested additional cognitive mechanisms and brain regions linked with reward processing. We propose that in complex reward contexts, valuation may be influenced by emotional factors, probabilistic external factors, and/or other subjective internal factors. The behavior profiles for MAGs 3-7 included terms representing cognitive mechanisms involved in processing a variety of the peripheral aspects of reward contingencies that often influence (augment or discount) valuation. Based on the results of our functional decoding analyses, we interpreted the functional dissociation of MAGs-3-7 to reflect the processing of differential value modulators. Depending on the type of task being employed, value could be modulated by internally focused processes (e.g., emotion, physiological/affective states, episodic memory of prior experiences, and personal beliefs or preferences), and/or externally focused processes (e.g., performance monitoring, executive attention, risk and probability calculation).

### Emotional influences on valuation

MAG-3 displayed convergent activity in the amygdala, hippocampus, left posterior insula, and SMA and was associated with affect-related terms such as *reactivity*, *emotion*, *emotional stimuli*, *anxiety*, *fearful*, and *happy.* Reward-related contexts often engender acute emotional states necessitating emotion identification and/or regulation (Quartz, 2009; Seymour & Dolan, 2008). Affective states such as fear, stress, anger, and excitement, and the capacity to regulate these emotions, impact reward processing and goal-directed decision-making (Coricelli, Dolan & Sirigu, 2007; Lawrence, Allen & Chanen, 2010; Paulus & Yu, 2012). *External* reward contexts may involve emotional or arousing stimuli (e.g., *happy* or *angry* facial *expressions*; deprecating or praising feedback), which in turn may evoke internal emotional reactions (*reactivity*, *anxiety*, *fearful*, *emotion*) or require *internal* processes (*emotion regulation*). Therefore, in the context of our meta-analytic corpus, it is plausible that convergent activity patterns linked with emotional processing would not discreetly cluster into an internal or external subgrouping of studies but rather, may represent a distinct neurocomputational influence on reward processing as observed in MAG-3.

### External influences on valuation

Both MAG-4 and MAG-5 exhibited convergent activity in the IPS and LPFC, major hubs of the ECN (Seeley et al., 2007). The ECN is thought to underlie “task-on attention” when tasks involve executive attention, working memory, inhibition, cognitive flexibility, and planning (Brown et al., 2019; Niendam et al., 2012) and has been implicated in reward-motivated cognitive control and decision-making (Hobkirk et al., 2019; Krmpotich, 2013; Tanji & Hoshi, 2008). Additionally, the IPS and LPFC have been associated with cue-based attention orienting (Anderson, 2017) and monitoring peripheral aspects of reward tasks such as stimuli presentation patterns and a stimulus’s associated probability or risk levels (Blankenstein et al., 2018; Glascher et al., 2010; Guo et al., 2018; Huettel, Mack & McCarthy, 2002; Rushworth & Behrens, 2008; Sela, Kilim & Lavidor, 2012). The correspondence between MAG-4 and MAG-5 and the ECN is intuitive considering the terms associated with these MAGs suggest a role in processing external aspects of the task context (MAG-4: *monitoring*, *difficulty*, *decision*, *conflict*; MAG-5: *performance*, *calculation*, *interference, arithmetic*), employing executive control (MAG-4: *cognitive control*, MAG-5: *executive*, *attentional*), and maintaining information in working memory (MAG-4 & 5: *working memory*, *maintenance*, *load*).

Notably, MAG-4 also displayed robust convergent activity in core nodes of the SN (Seeley et al., 2007) including the bilateral anterior insulae, dACC, and thalamic nuclei, which were not observed in MAG-5. The SN is another large-scale brain network thought to be involved in detecting salient stimuli and accordingly orienting attentional resources (Menon & Uddin, 2010; Sridharan, Levitin & Menon, 2008; Sutherland et al., 2012). Additionally, MAG-4 included two bilateral clusters centered over the habenular nuclei that extended into the thalamus and midbrain. MAG-4’s pattern of convergent activation also resembled previous accounts of human habenula connectivity (Ely et al., 2019; Ely et al., 2016) particularly during error processing (Ide & Li, 2011). The habenula has been implicated in processing negative outcomes and error monitoring (Flannery et al., 2019) (Baker et al., 2016; Li et al., 2008; Ullsperger & Cramon, 2003) and is of emerging interest in the context of reward processing and pathologies characterized by disruptions thereof, such as addiction and depression (Batalla et al., 2017; Flannery et al., 2019; Loonen & Ivanova, 2016; Mathis & Kenny, 2018). Thus, we found it particularly noteworthy that MAG-4 was uniquely associated with terms suggesting a potential role in performance monitoring (*difficulty*, *stop*, *conflict*, *monitoring*, *cognitive control*). In line with the bottom-up, salience detecting, and attention-orienting roles of the SN, preclinical research has demonstrated a habenula-ACC circuit in which errors are detected in the habenula and maintained in the ACC (Kawai et al., 2015). Then, through interactions with the ACC, the anterior insula is thought to use this information to appropriately mediate dynamic switching between DMN and ECN activity (Kerns et al., 2004; Menon & Uddin, 2010; Sridharan, Levitin & Menon, 2008; Sutherland et al., 2012).

Overall our functional decoding of MAG-4 and MAG-5 suggest that they may be involved in executive functions needed during reward-processing tasks, such as monitoring task performance or task difficulty, assessing associated risk levels, or calculating the probabilistic nature of reward contingencies. Further, the dissociated convergent activity of these two MAGs may highlight discrete bottom-up, cognitive control (MAG-4) and higher-order, calculation-based (MAG-5) executive functions during reward processing.

### Internal influences on valuation

In addition to external influences, valuation could be influenced by a variety of internal factors specific to an individual’s current hedonic state and past experiences; both of which are modulated by physiological, social, and/or cultural influences (Bartra, McGuire & Kable, 2013; McClure et al., 2004; Rangel, 2008; Yoder & Decety, 2018). Both MAG-6 and MAG-7 were associated with terms suggesting such an internally-oriented focus such that MAG-6’s behavior profile was indicative of self-referential abstract mentalizing (*self-referential*, *mentalizing*, *mental states*, *beliefs*, *moral*) and MAG-7’s profile was indicative of using personal preferences and prior experience to form subjective values (*value*, *valence*, *arousal*, *autobiographical*, *emotion*, *neutral, referential*).

Both MAGs were characterized by topologic resemblance to the DMN which further supported such an interpretation, given the known link between DMN regions and internally focused tasks involving theory of mind, autobiographical memory retrieval, and other forms of self-referential thought (Hamilton et al., 2015; Mars et al., 2012; Philippi et al., 2015; Spreng & Grady, 2009); (Qin & Northoff, 2011). However, despite some similarities, dissociable areas of activity convergence were noteworthy between these MAGs, such that MAG-6 displayed convergence in more dorsal regions of the mPFC and PCC, whereas MAG-7 displayed more ventral convergence in the mPFC, medial OFC, and PCC. Further, MAG-6 included convergent activity in the TPJ, precuneus, lateral inferior frontal gyrus, and middle temporal gyrus that was not seen in MAG-7. Instead, MAG-7 displayed convergent activity in a cluster encompassing the right amygdala and hippocampus.

Regions included specifically in MAG-6 have been broadly associated with tasks evoking theory of mind and self-referential thought (Mars et al., 2012; Qin & Northoff, 2011). Additionally, MAG-7 includes brain regions (vmPFC, medial OFC, amygdala, PCC) that have been implicated in subjective valuation networks defined by previous work (Bartra, McGuire & Kable, 2013; Levy & Glimcher, 2012; Peters & Buchel, 2010). For example, Acikalin and colleagues (2017) demonstrated a meta-analytic, functional overlap between the DMN and what they termed the subjective valuation network (SVN). Key nodes displaying DMN and SVN overlap were also reflected in MAG-7 of our meta-analytic *k*-means clustering analysis. Overall, our functional decoding of MAG-6 and MAG-7 demonstrated how two aspects of internally focused attention may be functionally dissociated into similar networks that differ in their recruitment of the mPFC (ventral or dorsal) and other regions (right amygdala or TPJ). The dissociated convergent activity across these two MAGs corresponds with previous work exploring the multifaceted functions associated with the DMN (Acikalin, Gorgolewski & Poldrack, 2017; Mars et al., 2012; Qin & Northoff, 2011).

### Comprehensive meta-analysis: Integration of understudied functions

The inclusion criteria for our corpus of experiments encompassed a wider range of reward-related tasks than previous meta-analyses have considered. This allowed us to provide a unique meta-analytic account of brain regions that are not as commonly linked to reward processing, but are nonetheless, vital for its realization. Previous neuroimaging meta-analyses of reward processing have focused on specific aspects such as food processing (van der Laan et al., 2011) and subjective valuation (Bartra, McGuire & Kable, 2013), or specific comparisons such as passive anticipation versus consumption of rewards (Diekhof et al., 2012) and receipt of primary versus secondary rewards (Sescousse et al., 2013). While our sample of reward processing results was constrained to the perhaps disproportionate frequency distribution of certain topics and tasks characterized in the extant neuroimaging literature, our sample did not exclude experimental contrasts based on outcome valance (i.e., positive, negative), outcome type (i.e., monetary, food, verbal), or processing stage (i.e., decision, anticipation, outcome delivery). Additionally, while existing meta-analyses manually classified datasets into aspects of interest to subsequently characterize associated brain activation patterns (e.g., Liu, 2011), our meta-analytic, data-driven parsing of experimental contrasts allowed for groups of activity patterns to emerge organically and unbiased by prior assumptions.

### Methodological considerations

Our clustering results may be influenced by characteristics of the *k*-means clustering algorithm, specifically its sensitivity to spherical clusters and its assumption that input data are linearly separable (Bottenhorn et al., 2019). Further, while this approach is, for the most part, data-driven, it requires specification of certain parameters. Specifically, we chose 1,000 as the number of iterations performed to ensure that each solution minimized the point-to-centroid distance. This decision was made based on prior work (Bottenhorn et al., 2019; Kanungo, 2004). We also selected the additive inverse of the Pearson’s correlation coefficient as the metric for calculating pair-wise distances between variables as it has been previously recommended to maximize distances between spatial patterns of activity maps (Laird et al., 2015). Exploring the impact of these parameters on clustering results is beyond the scope of the current work.

While we used metrics to determine viable clustering solutions, the optimal solution is not always an obvious selection. Our metrics identified three viable model orders and we ultimately selected the solution that identified the greatest number of neurocognitively plausible MAGs. However, we also report assessments of the two other viable solutions in the supplemental material and believe that examination of all three viable model orders provides increased transparency regarding the integration and segregation of our data across degrees of meta-analytic, *k*-means clustering parcellation. Additionally, as with all meta-analyses, our results are potentially influenced by any reporting biases present in the extant literature. Finally, while our functional decoding yielded outcomes reflective of the broader literature, more corpus-specific and domain-specific insights might have been achieved with a functional decoding approach based on annotation of experiments in our corpus (see supplemental materials for further discussion on manual corpus-specific versus automated generalized annotation strategies). Approaches allowing for unbiased, objective, and yet detailed annotation of corpus-specific experiments may benefit future research.

### Conclusion

We dissociated meta-analytic groupings of reward processing experiments based on the similarity between each experiment’s pattern of activation using a meta-analytic *k*-means clustering approach. The resulting seven meta-analytic groupings represented patterns of activity consistently occurring across reward processing tasks. Our functional decoding analyses revealed that these dissociable brain activity patterns could be mapped onto discrete mental processes within a heuristic framework of valuation. Specifically, we observed two frontal-striatal pathway MAGs, one displaying convergent activity in a VS-vmPFC network (MAG-1) and another displaying convergent activity in a DS-dmPFC network (MAG-2). While our functional decoding linked both MAGs to anticipating outcomes and encoding RPEs, these MAGs appeared to be consistent with specialized roles described in actor-critic models. Further, we identified five additional MAGs that correspond to recognized limbic (MAG-3), SN (MAG-4), ECN (MAG-5), DMN (MAG-6), and SVN (MAG-7) activity patterns. Functional decoding linked these MAGs to processing distinct emotional, internal, and external influences present across complex reward contexts. These findings demonstrate the extensive variety of activity patterns involved in aspects of valuation and highlight the role of commonly observed large-scale brain networks in certain aspects of reward processing tasks.

## ACKNOWLEDGEMENTS

Primary support for this project was provided by NIH R01DA041353 (MTS, ARL, MCR, RP) and NIH U54MD012393 (JSF, MTS). Contributions from co-authors were also provided with support from NSF 1631325 (ARL, MCR, TS), NIH K01DA037819 (MTS), NIH U01DA041156 (ARL, MTS, MCR, KLB, LDHB), NSF REAL DRL-1420627 (ARL), and NSF CNS 1532061 (ARL). We thank the FIU Instructional & Research Computing Center (IRCC, http://ircc.fiu.edu) for providing access to the HPC computing resources that contributed to the generation of the research results reported within this paper.

## OPEN PRACTICES STATEMENT

The authors have released all code and data associated with this manuscript. The code and tabular data are available on GitHub (https://github.com/NBCLab/reward-processing-meta-analysis), and the unthresholded maps of each MAG are available on NeuroVault (https://neurovault.org/collections/5070/).

