## Supplemental Information for "Meta-analytic clustering dissociates brain activity and behavior profiles across reward processing paradigms"

### SUPPLEMENTAL CONTENT


- Table S1. Published articles included in the current reward processing corpus
- Table S2. BrainMap paradigm class composition of the corpus
- Table S3. Convergent activity coordinates for each meta-analytic grouping (*k*=7 model order)
- ADDITIONAL VIABLE CLUSTERING SOLUTIONS: *POST HOC* COMPARISON
    - Text. *k* = 5 solution in relation to *k* = 7 solution
    - Figure S1. Brain activity profiles for the *k* = 5 solution
    - Figure S2. Behavioral profiles for the *k* = 5 solution
    - Table S4. NeuroSynth functional decoding results for the *k* = 5 solution
    - Text. *k* = 4 solution in relation to *k* = 5 solution
    - Figure S3. Brain activity profiles for the *k* = 4 solution
    - Figure S4. Behavioral profiles for the *k* = 4 solution
    - Table S5. NeuroSynth functional decoding results for the *k* = 4 solution
- Figure S5. *K*-means clustering cross-correlation matrix (*k* = 7 model order)
- Figure S6. Comparison of convergent activity from MAG-1 vs. -2 and from MAG-6 vs. -7
- MANUAL (vs. automated) ANNOTATION FOR FUNCTIONAL DECODING
    - Text. Corpus-specific manual annotation of experiments (*k* = 5 solution): Rationale
    - Table S6. Term glossary and frequency distribution across MAGs
    - Text. Corpus-specific manual functional decoding (*k* = 5 solution): Methods
    - Text. Corpus-specific manual functional decoding (*k* = 5 solution): Results
    - Figure S7. Behavior profiles derived from corpus-specific manual decoding
    - Figure S8. Effect size of term-MAG associations (*k* = 5 solution)
- CORPUS ARTICLES REFERENCES
- SUPPELMENTAL REFERENCES

**Table S1. Published articles included in the current reward processing corpus.** 176 studies contributed experimental contrasts (minimum = 1, maximum = 23) to the corpus.

| | PubMed ID | Authors | Year | # of contrasts |
|---|---|---|---|---|
| 1 | 17265148 | Abler, Erk & Walter | 2007 | 3 |
| 2 | 16487726 | Abler et al. | 2006 | 2 |
| 3 | 16675403 | Adcock et al. | 2006 | 12 |
| 4 | 12948722 | Akitsuki et al. | 2003 | 8 |
| 5 | 19961940 | Alexander & Brown | 2010 | 4 |
| 6 | 19071223 | Ballard & Knutson | 2009 | 3 |
| 7 | 22336565 | Balodis et al. | 2012 | 12 |
| 8 | 19560123 | Beck et al. | 2009 | 4 |
| 9 | 17676057 | Behrens et al. | 2007 | 2 |
| 10 | 19587291 | Bickel et al. | 2009 | 9 |
| 11 | 17140674 | Bjork & Hommer | 2007 | 9 |
| 12 | 14985419 | Bjork et al. | 2004 | 4 |
| 13 | 18851716 | Bjork, Knutson & Hommer | 2008 | 4 |
| 14 | 17079666 | Blair et al. | 2006 | 9 |
| 15 | 15907305 | Bolla et al. | 2005 | 1 |
| 16 | 15142963 | Bolla et al. | 2004 | 2 |
| 17 | 19524531 | Boorman et al. | 2009 | 2 |
| 18 | 18509047 | Bray et al. | 2008 | 1 |
| 19 | 11395019 | Breiter et al. | 2001 | 4 |
| 20 | 10706432 | Brunia et al. | 2000 | 2 |
| 21 | 17188518 | Budhani et al. | 2007 | 2 |
| 22 | 22338036 | Burger & Stice | 2012 | 2 |
| 23 | 20589242 | Burke et al. | 2010 | 3 |
| 24 | 19242558 | Camara, Rodriguez-Fornells & Munte | 2008 | 6 |
| 25 | 18042401 | Chandrasekhar et al. | 2008 | 4 |
| 26 | 19793990 | Chib et al. | 2009 | 4 |
| 27 | 19812332 | Christopoulos et al. | 2009 | 2 |
| 28 | 19217383 | Clark et al. | 2009 | 3 |
| 29 | 17997112 | Cohen, Elger & Weber | 2008 | 5 |
| 30 | 19843618 | Cooper et al. | 2009 | 9 |
| 31 | 17904386 | Cooper & Knutson | 2008 | 3 |
| 32 | 16116457 | Coricelli et al. | 2005 | 12 |
| 33 | 15758183 | Cox, Andrade & Johnsrude | 2005 | 1 |
| 34 | 11239442 | Critchley, Mathias & Dolan | 2001 | 3 |

| | | | | |
|---|---|---|---|---|
| 35 | 18309087 | D'Ardenne et al. | 2008 | 1 |
| 36 | 16778890 | Daw et al. | 2006 | 4 |
| 37 | 11110834 | Delgado et al. | 2000 | 3 |
| 38 | 17850241 | Dillon et al. | 2008 | 7 |
| 39 | 16033924 | Dreher, Kohn & Berman | 2006 | 6 |
| 40 | 21612768 | Duka et al. | 2011 | 4 |
| 41 | 18445214 | Elliott, Agnew & Deakin | 2008 | 2 |
| 42 | 10934265 | Elliott, Friston & Dolan | 2000 | 5 |
| 43 | 9347486 | Elliott, Frith & Dolan | 1997 | 6 |
| 44 | 12514228 | Elliott et al. | 2003 | 3 |
| 45 | 15006665 | Elliott et al. | 2004 | 3 |
| 46 | 10215087 | Elliott, Rees & Dolan | 1999 | 4 |
| 47 | 9626713 | Elliott et al. | 1998 | 3 |
| 48 | 19640506 | Elman et al. | 2009 | 1 |
| 49 | 19308261 | Engelmann et al. | 2009 | 7 |
| 50 | 19576868 | Engelmann & Tamir | 2009 | 5 |
| 51 | 15850746 | Ernst et al. | 2005 | 10 |
| 52 | 15327927 | Ernst et al. | 2004 | 3 |
| 53 | 19047075 | Ersner-Hershfield, Wimmer & Knutson | 2009 | 7 |
| 54 | 18985124 | Feinstein, Stein & Paulus | 2006 | 2 |
| 55 | 18793731 | Finger et al. | 2008 | 8 |
| 56 | 20357071 | Fleming et al. | 2010 | 7 |
| 57 | 17656073 | Frangou et al. | 2008 | 1 |
| 58 | 19783412 | Freyer et al. | 2009 | 2 |
| 59 | 15588617 | Fukui et al. | 2005 | 1 |
| 60 | 16682235 | Fukui et al. | 2006 | 1 |
| 61 | 17286837 | Galvan et al. | 2007 | 2 |
| 62 | 18550593 | Glascher, Hampton & O'Doherty | 2009 | 7 |
| 63 | 17202543 | Goldstein et al. | 2007 | 1 |
| 64 | 20600994 | Guitart-Masip, Talmi & Dolan | 2010 | 1 |
| 65 | 17698008 | Hampton et al. | 2007 | 5 |
| 66 | 16899731 | Hampton, Bossaerts & O'Doherty | 2006 | 5 |
| 67 | 18427116 | Hampton, Bossaerts & O'Doherty | 2008 | 4 |
| 68 | 20357127 | Han et al. | 2010 | 1 |
| 69 | 20071521 | Hare et al. | 2010 | 5 |
| 70 | 19407204 | Hare, Camerer & Rangel | 2009 | 4 |

| | | | | |
|---|---|---|---|---|
| 71 | 18509023 | Hare et al. | 2008 | 3 |
| 72 | 20105435 | Hartstra et al. | 2010 | 5 |
| 73 | 14973239 | Haruno et al. | 2004 | 4 |
| 74 | 16339445 | Hsu et al. | 2005 | 6 |
| 75 | 19228976 | Hsu et al. | 2009 | 2 |
| 76 | 17007234 | Huettel | 2006 | 4 |
| 77 | 16504951 | Huettel et al., | 2006 | 1 |
| 78 | 18439412 | Izuma, Saito & Sadato | 2008 | 4 |
| 79 | 19515916 | Jocham et al. | 2009 | 4 |
| 80 | 16139525 | Juckel et al. | 2006 | 2 |
| 81 | 20510371 | Kahnt et al. | 2011 | 2 |
| 82 | 20231475 | Kahnt et al. | 2010 | 2 |
| 83 | 16802856 | Kim, Shimojo & O'Doherty | 2006 | 8 |
| 84 | 14568478 | Kirsch et al. | 2003 | 4 |
| 85 | 11459880 | Knutson et al. | 2001 | 5 |
| 86 | 17916330 | Knutson et al. | 2008 | 4 |
| 87 | 15260961 | Knutson et al. | 2004 | 4 |
| 88 | 11726774 | Knutson et al. | 2001 | 4 |
| 89 | 12595181 | Knutson et al. | 2003 | 3 |
| 90 | 17196537 | Knutson et al. | 2007 | 3 |
| 91 | 15888656 | Knutson et al. | 2005 | 8 |
| 92 | 10875899 | Knutson et al. | 2000 | 2 |
| 93 | 18388729 | Knutson et al. | 2008 | 3 |
| 94 | 18549791 | Knutson et al. | 2008 | 23 |
| 95 | 19032746 | Koeneke et al. | 2008 | 9 |
| 96 | 17765572 | Kramer et al. | 2007 | 7 |
| 97 | 16129404 | Kuhnen & Knutson | 2005 | 4 |
| 98 | 18320179 | Labudda et al. | 2008 | 2 |
| 99 | 18787233 | Lawrence et al. | 2009 | 5 |
| 100 | 19015090 | Lee et al. | 2008 | 2 |
| 101 | 19770058 | Linke et al. | 2010 | 5 |
| 102 | 16426719 | Little et al. | 2006 | 7 |
| 103 | 17460071 | Liu et al. | 2007 | 6 |
| 104 | 17712267 | Marco-Pallares, Muller & Munte | 2007 | 1 |
| 105 | 17292631 | Marsh et al. | 2007 | 2 |
| 106 | 11703464 | Martin-Soelch et al. | 2001 | 3 |
| 107 | 12911764 | Martin-Soelch et al. | 2003 | 1 |
| 108 | 11545466 | Martin-Solch et al. | 2001 | 3 |

| | | | | |
|---|---|---|---|---|
| 109 | 15486494 | Matthews et al. | 2004 | 2 |
| 110 | 12718866 | McClure, Berns & Montague | 2003 | 2 |
| 111 | 15486304 | McClure et al. | 2004 | 2 |
| 112 | 20138482 | Miedl et al. | 2010 | 2 |
| 113 | 19726640 | Mitchell et al. | 2009 | 2 |
| 114 | 17950474 | Mitchell et al. | 2008 | 1 |
| 115 | 17717184 | Mobbs et al. | 2007 | 17 |
| 116 | 15945130 | Nieuwenhuis et al. | 2005 | 5 |
| 117 | 15978024 | Nieuwenhuis et al. | 2005 | 1 |
| 118 | 15087550 | O'Doherty et al. | 2004 | 5 |
| 119 | 11135651 | O'Doherty et al. | 2001 | 9 |
| 120 | 19864559 | Palminteri et al. | 2009 | 8 |
| 121 | 11133312 | Paulus et al. | 2001 | 4 |
| 122 | 12948701 | Paulus et al. | 2003 | 2 |
| 123 | 16929307 | Pessiglione et al. | 2006 | 5 |
| 124 | 20016088 | Peters & Buchel | 2009 | 12 |
| 125 | 20399735 | Peters & Buchel | 2010 | 7 |
| 126 | 17855612 | Plassmann, O'Doherty & Rangel | 2007 | 6 |
| 127 | 11960021 | Pochon et al. | 2002 | 2 |
| 128 | 16880132 | Preuschoff, Bossaerts & Quartz | 2006 | 7 |
| 129 | 18337404 | Preuschoff, Quartz & Bossaerts | 2008 | 3 |
| 130 | 15528079 | Ramnani et al. | 2004 | 2 |
| 131 | 12571121 | Ramnani & Miall | 2003 | 4 |
| 132 | 18582578 | Rao et al. | 2008 | 7 |
| 133 | 15907318 | Remijnse et al. | 2005 | 8 |
| 134 | 17088503 | Remijnse et al. | 2006 | 3 |
| 135 | 15538191 | Rilling et al. | 2004 | 1 |
| 136 | 10516320 | Rogers et al. | 1999 | 8 |
| 137 | 17698371 | Sailer et al. | 2007 | 5 |
| 138 | 17468751 | Samanez-Larkin et al. | 2007 | 8 |
| 139 | 18399882 | Samanez-Larkin et al. | 2008 | 1 |
| 140 | 12805551 | Sanfey et al. | 2003 | 1 |
| 141 | 17655834 | Schaefer & Rotte | 2007 | 1 |
| 142 | 16950228 | Scheres et al. | 2007 | 1 |
| 143 | 18097655 | Schlagenhauf et al. | 2008 | 4 |
| 144 | 18032658 | Schonberg et al. | 2007 | 4 |
| 145 | 17475790 | Seymour et al. | 2007 | 4 |

| | | | | |
|---|---|---|---|---|
| 146 | 18947356 | Shamosh et al. | 2008 | 2 |
| 147 | 19718655 | Simoes-Franklin et al. | 2010 | 3 |
| 148 | 18804540 | Smith et al. | 2009 | 3 |
| 149 | 20164333 | Smith et al. | 2010 | 6 |
| 150 | 19174537 | Spreckelmeyer et al. | 2009 | 2 |
| 151 | 19521264 | Sripada et al. | 2009 | 1 |
| 152 | 17996464 | Strohle et al. | 2008 | 2 |
| 153 | 18579749 | Tanaka, Balleine & O'Doherty | 2008 | 2 |
| 154 | 9175118 | Thut et al. | 1997 | 1 |
| 155 | 18987206 | Tobler et al. | 2008 | 3 |
| 156 | 17122317 | Tobler et al. | 2007 | 3 |
| 157 | 19490086 | Tricomi, Balleine & O'Doherty | 2009 | 6 |
| 158 | 12764119 | Ullsperger & von Cramon | 2003 | 5 |
| 159 | 19793875 | Valentin & O'Doherty | 2009 | 4 |
| 160 | 16574168 | van Leijenhorst, Crone & Bunge | 2006 | 8 |
| 161 | 21389226 | Venkatraman et al. | 2011 | 5 |
| 162 | 19477159 | Venkatraman et al. | 2009 | 4 |
| 163 | 19349237 | Volkow et al. | 2009 | 1 |
| 164 | 12814578 | Volz, Schubotz & von Cramon | 2003 | 2 |
| 165 | 15006651 | Volz, Schubotz & von Cramon | 2004 | 3 |
| 166 | 19307555 | Weber et al. | 2009 | 3 |
| 167 | 18710652 | Weber & Huettel | 2008 | 5 |
| 168 | 17216152 | Wittmann, Leland & Paulus | 2007 | 2 |
| 169 | 17521924 | Wrase et al. | 2007 | 7 |
| 170 | 17291784 | Wrase et al. | 2007 | 2 |
| 171 | 19805082 | Wunderlich, Rangel & O'Doherty | 2009 | 6 |
| 172 | 19185567 | Xu et al. | 2009 | 5 |
| 173 | 18842669 | Xue et al. | 2009 | 6 |
| 174 | 16971537 | Yacubian et al. | 2006 | 11 |
| 175 | 20600178 | Zheng, Wang & Zhu | 2010 | 3 |
| 176 | 15134646 | Zink et al. | 2004 | 3 |

**<u>Table S2.</u> BrainMap paradigm class composition of the corpus.** Percentage of experiments in the corpus archived under each BrainMap paradigm class.

| Paradigm Class | Percentage of experiments in corpus |
|---|---|
| Reward | 94.9 |
| Task Switching | 6.4 |
| Delay Discounting | 5.6 |
| Go-NoGo | 2.9 |
| Visuospatial | 2.9 |
| Gambling | 2.7 |
| Wisconsin Card Sorting Test | 2.5 |
| Reasoning Problem Solving | 1.3 |
| Tower of London | 1.2 |
| Finger Tapping Button Press | 0.9 |
| Saccades | 0.8 |
| Taste | 0.8 |

<u>Note.</u> As studies included in the corpus could be archived under multiple paradigm classes, the percentages of corpus experiments in each paradigm class are not expected to sum to 100%.

**Table S3. Convergent activity coordinates for each meta-analytic grouping (*k* = 7 model order)**

| Meta-Analytic Grouping | Peak | Region | Volume ($mm^3$) | x | y | z |
|---|---|---|---|---|---|---|
| 1 | 1 | Left ventral striatum | 52944 | -12 | 10 | -6 |
| | | Right accumbens | | 12 | 10 | -6 |
| | | Right claustrum | | 34 | 22 | -8 |
| | | Right thalamus | | 2 | -14 | 10 |
| | 2 | Medial orbital frontal gyrus (BA 10) | 3168 | 0 | 50 | -10 |
| | | Left medial orbital frontal | | -12 | 42 | -14 |
| | 3 | Anterior cingulate (BA 32) | 4056 | -4 | 38 | 16 |
| | | Right middle dorsal cingulate | | 2 | 28 | 34 |
| | | Right middle dorsal cingulate | | 4 | 18 | 36 |
| | 4 | Posterior cingulate | | -2 | -32 | 32 |
| 2 | 1 | Left caudate | 39536 | -10 | 4 | 6 |
| | | Right caudate | | 12 | 4 | 8 |
| | | Right pallidum | | 14 | 4 | 0 |
| | | Left putamen | | -18 | 4 | 6 |
| | 2 | Dorsal medial frontal gyrus (BA 32) | 6248 | 0 | 10 | 46 |
| | | Right dorsal cingulate | | 6 | 22 | 30 |
| | | Dorsal medial frontal gyrus (BA 6) | | 0 | 0 | 56 |
| | | Left dorsal medial frontal gyrus (BA 24) | | -4 | -8 | 48 |
| | 3 | Right posterior cingulate (BA 23) | 1408 | 2 | -24 | 34 |
| 3 | 1 | Right amygdala | 8848 | 26 | -2 | -14 |
| | | Right hippocampus | | 32 | -22 | -22 |
| | 2 | Left amygdala | 7712 | -24 | -2 | -14 |
| | | Left Putamen | | -28 | -6 | 0 |
| | | Left culmen | | -8 | -30 | -16 |
| | | Left parahippocampus | | -20 | -16 | -18 |
| | 3 | Right dorsal medial frontal gyrus (BA 6) | 1984 | 6 | -4 | 46 |
| 4 | 1 | Right anterior insula (BA 13) | 11208 | 36 | 20 | -4 |
| | 2 | Left anterior insula (BA 13) | 9864 | -30 | 22 | -2 |
| | 3 | Medial frontal gyrus (BA 8) | 11008 | 6 | 22 | 46 |
| | | Left cingulate (BA 32) | | -8 | 24 | 32 |
| | | Medial frontal gyrus (BA 6) | | -8 | 8 | 46 |
| | 4 | Left superior parietal gyrus (BA 7) | 2160 | -32 | -54 | 46 |
| | 5 | Right thalamus (medial dorsal nucleus) | 5512 | 8 | -12 | 10 |

| | | | | | | |
|---|---|---|---|---|---|---|
| | | Left habenula | | -4 | -26 | -2 |
| | | Right thalamus (anterior nucleus) | | -8 | -18 | 10 |
| | | Left thalamus (ventral posterior nucleus) | | 14 | -30 | -6 |
| | 6 | Left dorsal middle frontal gyrus (BA 6) | 1200 | -26 | 0 | 52 |
| | 7 | Left declive | 1488 | -40 | -64 | -20 |
| | 8 | Right precentral gyrus (BA 9) | 1544 | 44 | 14 | 30 |
| | | Right precentral gyrus (BA 9) | | 48 | 14 | 44 |
| | 9 | Right inferior parietal gyrus (BA 40) | 1608 | 46 | -52 | 54 |
| | 10 | Right posterior cingulate (BA 23) | 1168 | 6 | -24 | 28 |
| | 11 | Left middle frontal gyrus (BA 9) | 1136 | -38 | 26 | 34 |
| | 12 | Right middle frontal gyrus (BA 46) | 1680 | 44 | 32 | 20 |
| | | Right middle frontal gyrus (BA 9) | | 38 | 30 | 34 |
| | | Right middle frontal gyrus (BA 9) | | 34 | 44 | 32 |
| **5** | 1 | Dorsal medial frontal gyrus (BA 6) | 6280 | -2 | 32 | 32 |
| | | Right dorsal medial frontal gyrus (BA 6) | | 8 | 22 | 48 |
| | 2 | Right superior frontal gyrus (BA 9) | 7536 | 44 | 42 | 22 |
| | | Right superior frontal gyrus (BA 8) | | 42 | 32 | 40 |
| | | Right middle frontal gyrus (BA 46) | | 48 | 22 | 20 |
| | 3 | Left precentral gyrus (BA 6) | 4304 | -40 | 2 | 32 |
| | | Left precentral gyrus (BA 9) | | -46 | 22 | 32 |
| | | Left middle frontal gyrus (BA 9) | | -56 | 6 | 34 |
| | 4 | Right inferior parietal gyrus (BA 40) | 4056 | 48 | -42 | 52 |
| | | Right inferior parietal gyrus (BA 40) | | 44 | -38 | 44 |
| | 5 | Right superior parietal gyrus (BA 7) | 2496 | 26 | -62 | 44 |
| | | Right superior parietal gyrus (BA 7) | | 32 | -60 | 52 |
| | 6 | Right superior frontal gyrus (BA 10) | 1824 | 30 | 52 | 0 |
| | 7 | Left inferior parietal gyrus (BA 40) | 1344 | -42 | -44 | 44 |
| | 8 | Left middle frontal gyrus (BA 6) | 1224 | -28 | 2 | 52 |
| | 9 | Left lateral middle frontal gyrus | 1256 | -40 | 32 | 24 |
| | 10 | Right middle frontal gyrus (BA 6) | 2864 | 32 | 8 | 54 |
| | 11 | Left superior parietal gyrus (BA 7) | 1208 | -28 | -58 | 44 |
| **6** | 1 | Left anterior medial frontal gyrus (BA 9) | 11128 | -4 | 50 | 22 |
| | | Left anterior medial frontal gyrus (BA 10) | | -4 | 56 | 10 |
| | | Right anterior cingulate (BA 24) | | 4 | 32 | 12 |
| | | Right medial frontal gyrus (BA 10) | | 12 | 48 | 0 |
| | 2 | Left temporal parietal junction (39) | 6328 | -42 | -76 | 34 |
| | | Left superior temporal gyrus (BA 39) | | -48 | -56 | 32 |
| | | Left supra-marginal gyrus (BA 39) | | -58 | -50 | 34 |

| | | | | | | |
|---|---|---|---|---|---|---|
| | | Left supra-marginal gyrus (BA 39) | | -54 | -56 | 44 |
| | 3 | Right inferior frontal gyrus (BA 47) | 1752 | 48 | 36 | -14 |
| | | Right inferior frontal gyrus (BA 45) | | 52 | 38 | -4 |
| | 4 | Right middle temporal gyrus (BA 21) | 1472 | 66 | -26 | -8 |
| | | Right middle temporal gyrus (BA 21) | | 60 | -36 | -4 |
| | 5 | Right temporal parietal junction (BA 40) | 832 | 54 | -42 | 26 |
| | 6 | Left precuneus (BA 31) | 1456 | -4 | -48 | 34 |
| | | Left precuneus (BA 31) | | -6 | -60 | 28 |
| | 7 | Left inferior frontal gyrus (BA 47) | 1104 | -56 | 34 | -6 |
| **7** | 1 | Medial orbital frontal gyrus (BA 32) | 27504 | 0 | 56 | -8 |
| | | Subgenual anterior cingulate (BA 11) | | 0 | 34 | -18 |
| | | Right frontal pole (BA 10) | | 6 | 68 | 0 |
| | 2 | Left parahippocampus (amygdala) | 2888 | 28 | -6 | -20 |
| | 3 | Left posterior cingulate (BA 30) | 1424 | -6 | -54 | 14 |
| | | Left posterior cingulate (BA 31) | | -8 | -56 | 24 |
| | 4 | Left middle frontal gyrus (BA 47) | 880 | -30 | 38 | -10 |

Note. Peak and sub-peak coordinates (LPI), spatial volume and anatomical labeling informed by Eickhoff-Zilles macro labels from N27 (MNI_ANAT space) and Talairach-Tournoux atlas labels for clusters comprising each MAG's thresholded ($p_{cluster\text{-}level} < 0.05$ [FWE-corrected]; $p_{voxel\text{-}level} < 0.001$) ALE image.

### ADDITIONAL VIABLE CLUSTERING SOLUTIONS: *POST HOC* COMPARISON.

While the $k = 7$ model order was identified as a viable solution and selected for presentation in the main text, the hierarchy index and variation of information metrics indicated that the $k = 5$ model order was also a viable clustering solution (**main text Fig. 2**). Further, the $k = 4$ model order met criteria for the average silhouette metric (**main text Fig. 2**), also suggesting a viable solution. As such, in a *post hoc* assessment, we explored the organization of MAG activity patterns across the $k = 5$ (**Fig. S1**) and $k = 4$ (**Fig. S3**) MAG solutions and performed automated functional decoding (**Fig. S2 & Fig. S4**) using a NeuroSynth approach. Results are discussed in relation to the $k = 7$ outcomes found in the main text.

***k* = 5 model order in relation to *k* = 7 model order.** We believe our examination of three viable clustering model orders provides additional information regarding the integration and segregation of functional brain activity and cognitive-behavioral constructs across varying levels of meta-analytic parcellation. We observed that two MAGs in the $k = 5$ solution (**MAG-4$^{5}$** and **MAG-5$^{5}$**) appeared to be decomposed into multiple separate MAGs in the $k = 7$ solution. Specifically, MAG-4$^{5}$ was decomposed into MAG-4$^{7}$ and MAG-5$^{7}$ in the $k = 7$ solution and MAG-5$^{5}$ was decomposed into MAG-6$^{7}$ and MAG-7$^{7}$ in the $k = 7$ solution. Details of these instances of further parsing of experiments into dissociable MAGS are provided below.

**MAG-4$^{5}$**. Convergent activity clusters observed in MAG-4 of the $k = 5$ model order (MAG-4$^{5}$) (**Fig. S1, purple**) noted in the lateral prefrontal cortex (LPFC), intraparietal sulcus (IPS) and pre-supplementary motor area (pre-SMA) were observed in both MAG-4$^{7}$ (main text **Fig. 3, pink)** and MAG-5$^{7}$ (main text **Fig. 3, purple).** However, certain clusters observed in MAG-4$^{5}$ in the superior frontal cortex and parietal cortex were only observed in MAG-5$^{7}$ and convergent activity in the bilateral anterior insula and posterior cingulate cortex (PCC) were only observed in MAG-

4$^7$. Functional decoding also suggested that MAG-4$^5$ was decomposed into MAG-4$^7$ and MAG-5$^7$ in the $k$ = 7 solution, as MAG-4$^5$ included many of the same terms relating to working memory (*working memory*, *load, maintenance*), performance monitoring (*difficulty*, *performance*, *conflict*), calculating (*calculation*), and more general executive control (*cognitive control*, *task*; **Fig. S3, Table S4**), that were observed across both MAG-4$^7$ and MAG-5$^7$ in the $k$ = 7 solution (main text **Fig. 4**).

**MAG-5$^5$**. Similarly, the convergent activity from MAG-5 in the $k$ = 5 model order (MAG-5$^5$) was decomposed into two separate MAGs in the $k$ = 7 solution. In the $k$ = 5 solution, MAG-5$^5$ (**Fig. S1, blue**) displayed convergence in the ventral medial prefrontal cortex (vmPFC) and central medial prefrontal cortex (cmPFC), medial orbital frontal cortex (OFC), left angular gyrus, right amygdala, right hippocampus, PCC, temporal parietal junction (TPJ), and precuneus. However, in the $k$ = 7 solution (main text **Fig. 3**), MAG-5$^5$ activity was represented in two MAGs, one with convergent activity in the anterior cmPFC, TPJ, PCC, and right middle temporal lobe (MAG-6$^7$) and one with convergent activity in the vmPFC, mOFC, right amygdala, and more ventral PCC (MAG-7$^7$). The functional decoding of MAG-5$^5$ included terms that suggested both interpersonal and intrapersonal reflection (*mentalizing, theory mind, self-referential, autobiographical, social*) as well as terms that suggested subjective valuation (*value, valence, emotional, reward*; **Fig. S2, Table S4**). While the functional decoding of MAG-6$^7$ and MAG-7$^7$ both included terms suggesting general internal processing (*default, social, referential, autobiographical*; **Fig. 4, Table 3**), MAG-6$^7$ separately included terms (*theory of mind, self-referential, mental states, mentalizing, beliefs, and moral*) indicative of interpersonal and intrapersonal reflection, whereas MAG-7$^7$ separately included terms (*value, valence, emotion, neutral and arousal*) indicative of subjective value judgments. These functional decoding results suggest that the further parsing of experiments in the

$k = 7$ solution decomposed MAG-5[5] into two separate MAGs both with a potential role in internally focused attention yet one specializing in in abstract mentalizing (MAG-6[7]) and the other in valuation (MAG-7[7]). Stated from a different perspective, at a lower model order *(k* = 5) these two distinct yet related activity patterns and associated cognitive constructs were likely integrated in the same MAG (MAG-5[5]).

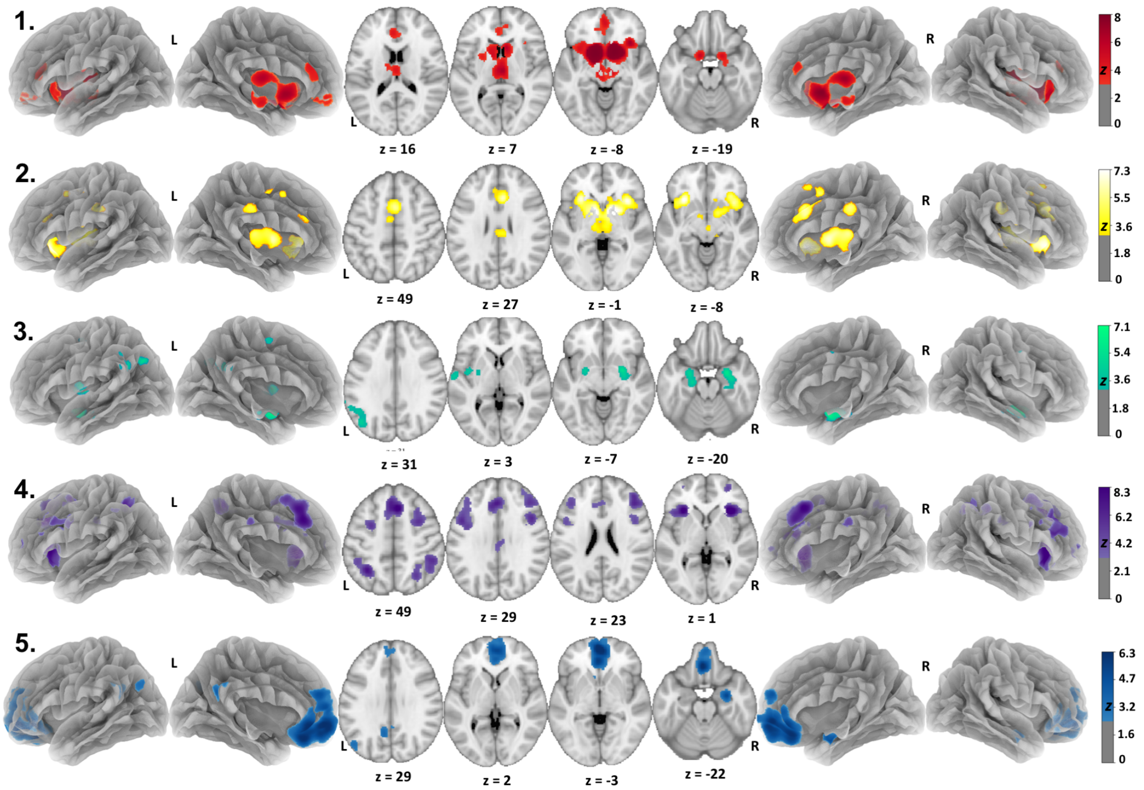


**Figure S1. Brain activity profiles associated with each meta-analytic grouping (MAG) of reward processing experiments (*k* = 5 model order).** ALE images identified significant ($p_{cluster\text{-}corrected} < 0.05$; $p_{voxel\text{-}level} < 0.001$) convergence in dissociable and distributed brain regions across each MAG. Unthresholded maps of each MAG are available on NeuroVault (https://neurovault.org/collections/5070/).

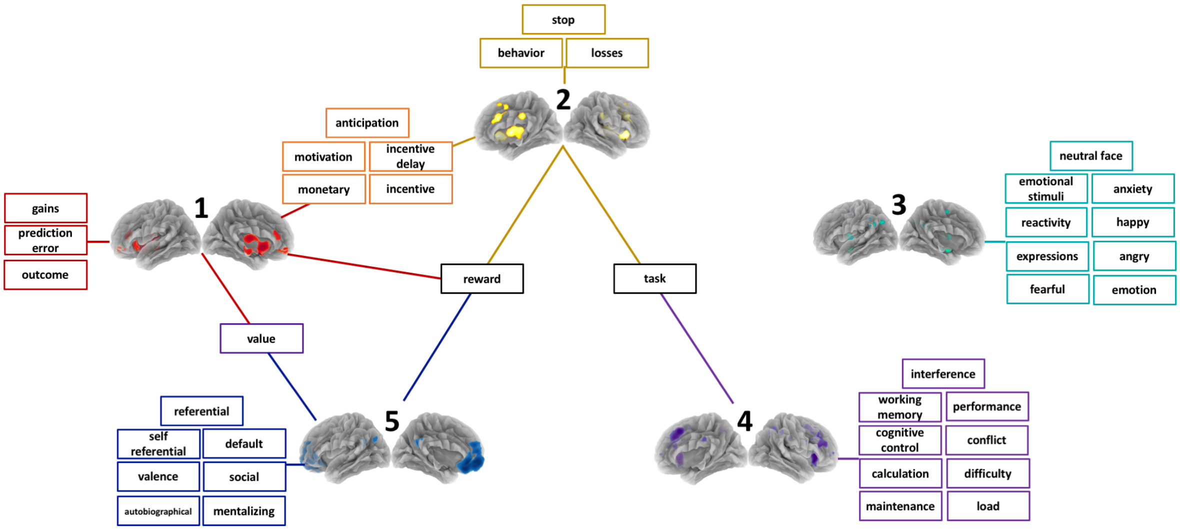


**<u>Figure S2.</u> Behavior profiles associated with each meta-analytic grouping (MAG) of reward processing experiments (*k* = 5 model order).** Behavior profiles consisted of NeuroSynth (NS) terms with the top 10 highest correlation values for each MAG representing the similarity between the MAG and activity patterns reported for terms in the NeuroSynth database (excluding anatomical terms). Lines connect MAGs to the terms making-up their unique behavior profile. Additionally, some terms or groups of terms are connected to multiple MAGs indicting that these terms had correlation values in multiple MAGs' top 10.

**Table S4. NeuroSynth (NS) terms composing each MAG's behavior profile and their respective correlation values (*k* = 5 model order).**

| MAG-1 | | MAG-2 | | MAG-3 | | MAG-4 | | MAG-5 | |
|---|---|---|---|---|---|---|---|---|---|
| **NS term** | ***r*** | **NS term** | ***r*** | **NS term** | ***r*** | **NS term** | ***r*** | **NS term** | ***r*** |
| monetary (2)[a] | 0.780 | anticipation (2)[g] | 0.321 | reactivity | 0.302 | task (2)[m] | 0.582 | default (2)[q] | 0.372 |
| reward (2)[b] | 0.766 | incentive (2)[h] | 0.289 | fearful (2)[k] | 0.301 | working memory (5)[n] | 0.510 | social | 0.305 |
| incentive (2)[c] | 0.754 | incentive delay | 0.288 | neutral | 0.292 | load (2)[o] | 0.382 | autobiographical (2)[r] | 0.295 |
| anticipation (2)[d] | 0.735 | monetary | 0.286 | emotion (3)[12] | 0.279 | difficulty (2)[p] | 0.326 | value | 0.262 |
| incentive delay | 0.701 | reward (3)[i] | 0.285 | happy | 0.266 | maintenance | 0.253 | referential | 0.254 |
| motivation (2)[e] | 0.677 | task | 0.219 | anxiety | 0.266 | cognitive control | 0.253 | valence | 0.246 |
| gains | 0.631 | motivation (2)[j] | 0.206 | expressions (2)[l] | 0.261 | performance | 0.241 | self referential | 0.232 |
| outcome (2)[f] | 0.616 | behavior | 0.206 | neutral faces | 0.259 | calculation | 0.241 | emotional | 0.229 |
| prediction error | 0.596 | stop | 0.193 | angry | 0.256 | interference | 0.238 | reward | 0.194 |
| value | 0.554 | losses | 0.188 | emotional stimuli | 0.268 | conflict | 0.236 | mentalizing | 0.191 |

Near Duplicates

| MAG-1 | MAG-2 | MAG-3 | MAG-4 | MAG-5 |
|---|---|---|---|---|
| [a] monetary reward | [g] reward anticipation | [k] fear | [n] tasks | [r] default mode |
| [b] rewards | [h] monetary incentive | [l] emotional, affective | [o] working, memory-wm, wm, memory | [s] autobiographical memory |
| [c] monetary incentive | [i] rewards, monetary reward | [m] facial expressions | [p] demands | |
| [d] reward anticipation | [j] motivational | | [q] task difficulty | |
| [e] motivational | | | | |
| [f] outcomes | | | | |

Note. Terms with the top 10 highest correlation values for each MAG representing the similarity between the MAG and activation patterns reported for terms automatically extracted from abstracts across functional neuroimaging studies archived in the NeuroSynth database (anatomical terms excluded). The number of near duplicates for any given term in a MAG's top 10 list is indicated in "()" following the term and the superscript labels the list of near duplicate terms in the lower section of the table.

***k* = 4 model order in relation to *k* = 5 model order.** Again, in the *k* = 4 lowest model order, we observed further condensing of MAGs that functional decoding accordingly linked to broader cognitive-behavior constructs. Specifically, subsets of convergent activity in MAG-1 (**Fig. S3, red**) of the *k* = 4 solution (MAG-$1^4$) appeared to be decomposed into two separate MAGs in the *k* = 5 solution (MAGs $1^5$ and $2^5$), indicating that the additional MAG included in the *k* = 5 solution segregated MAG-$1^4$ of the *k* = 4 solution into ventral and dorsal striatal-medial prefrontal cortex networks (**Fig. S1, red & yellow).** Further, NS terms associated with MAG-$1^4$ (*reward*, *anticipation*, *incentive delay*, *motivation, gains*; **Fig. S4, Table S5**) were associated with both MAG-$1^5$ and MAG-$2^5$ (**Table S4**). Overall, we suggest these outcomes indicated that, the *k* = 5 solution provided a more meaningful segregation of brain activity patterns during reward processing tasks that better captured a previously reported dissociation between ventral and dorsal frontal-striatal networks (see discussion section main text).

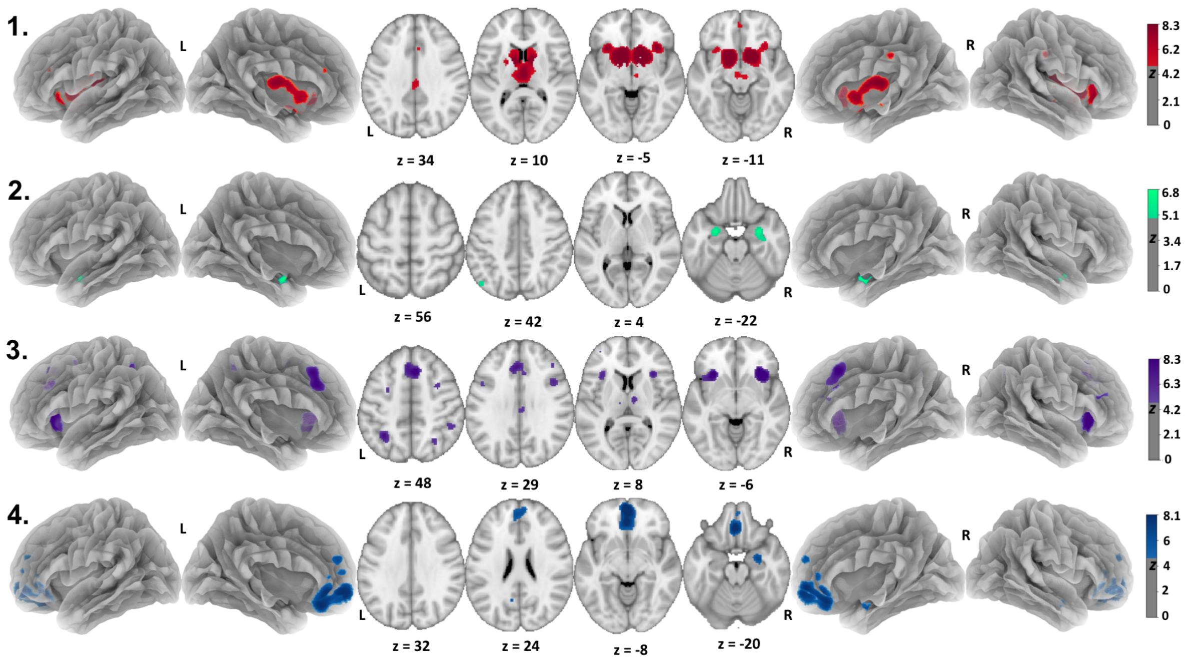


**Figure S3.** **Brain activity profiles associated with each meta-analytic grouping (MAG) of reward processing experiments ($k$ = 4 model order).** The figure depicts ALE images identifying significant ($p_{cluster\text{-}corrected}$ < 0.05; $p_{voxel\text{-}level}$ < 0.001) activity convergence for each MAG in the $k$ = 4 solution. Unthresholded maps of each MAG are available on NeuroVault (https://neurovault.org/collections/5070/).

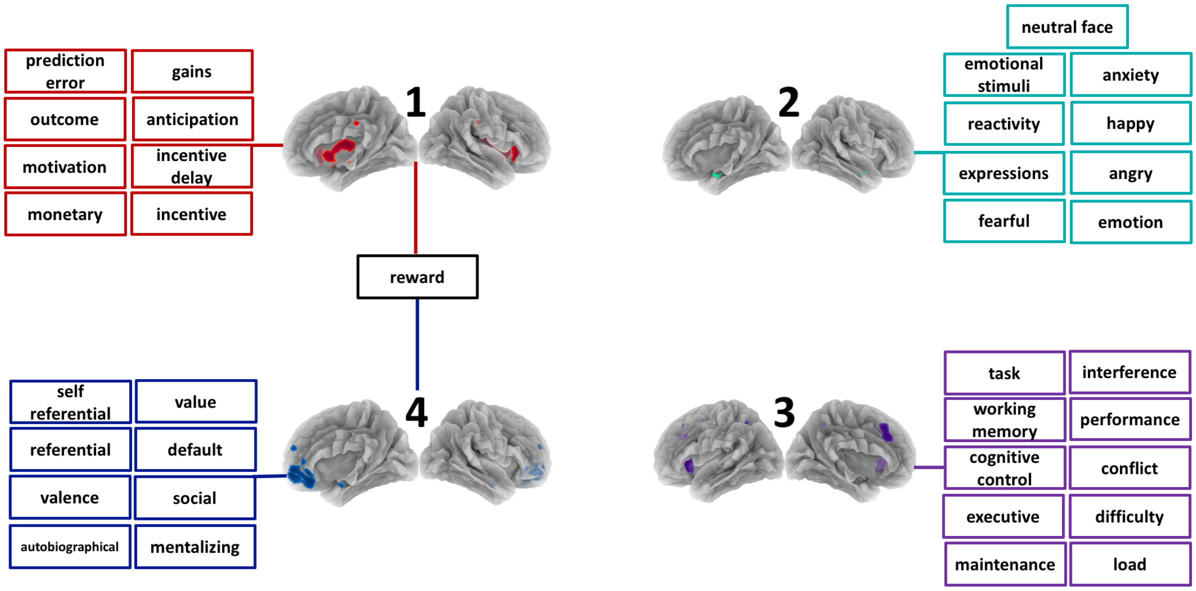


**Figure S4.** **Behavior profiles associated with each meta-analytic grouping (MAG) of reward processing experiments ($k$ = 4 model order).** Behavior profiles consisted of NeuroSynth (NS) terms with the top 10 highest correlation values for each MAG representing the similarity between the MAG and activity patterns reported for terms in the NeuroSynth database (excluding anatomical terms). Lines connect MAGs to the terms making-up their unique behavior profile. Additionally, some terms or groups of terms are connected to multiple MAGs indicting that these terms had correlation values in multiple MAGs' top 10.

**Table S5. NeuroSynth (NS) terms composing each MAG's behavior profile and their respective correlation values (*k* = 4 model order).**

| MAG-1 | | MAG-2 | | MAG-3 | | MAG-4 | |
|---|---|---|---|---|---|---|---|
| **NS term** | ***r*** | **NS term** | ***r*** | **NS term** | ***r*** | **NS term** | ***r*** |
| monetary | 0.731 | reactivity | 0.303 | task (2)[i] | 0.563 | default (2)[m] | 0.373 |
| reward (3)[a] | 0.717 | fearful (2)[e] | 0.301 | working memory (4)[j] | 0.467 | social | 0.307 |
| incentive (2)[b] | 0.712 | neutral (2)[f] | 0.292 | load (2)[k] | 0.353 | autobiographical (2)[n] | 0.291 |
| anticipation (2)[c] | 0.704 | emotion (3)[g] | 0.279 | difficulty (2)[l] | 0.317 | value | 0.264 |
| incentive delay | 0.669 | happy | 0.264 | conflict | 0.252 | referential | 0.249 |
| motivation (2)[d] | 0.617 | emotional stimuli | 0.264 | cognitive control | 0.246 | valence | 0.246 |
| gains | 0.572 | anxiety | 0.263 | maintenance | 0.242 | self referential | 0.228 |
| outcome | 0.538 | expressions (2)[h] | 0.257 | interference | 0.231 | emotional | 0.227 |
| prediction error | 0.513 | angry | 0.254 | performance | 0.230 | mentalizing | 0.196 |
| losses | 0.501 | pictures | 0.245 | executive | 0.221 | reward | 0.196 |

Near Duplicates

| | | | |
|---|---|---|---|
| [a] rewards, monetary reward | [e] fear | [i] tasks | [m] default mode |
| [b] monetary incentive | [f] neutral faces | [j] working, memory wm, memory | [n] autobiographical memory |
| [c] reward anticipation | [g] affective, emotional | [k] demands | |
| [d] motivational | [h] facial expressions | [l] task difficulty | |

Note. Terms with the top 10 highest correlation values for each MAG. The number of near duplicates for any given term in a MAG's top 10 list is indicated in "()" following the term and the superscript labels the list of near duplicate terms in the lower section of the table.

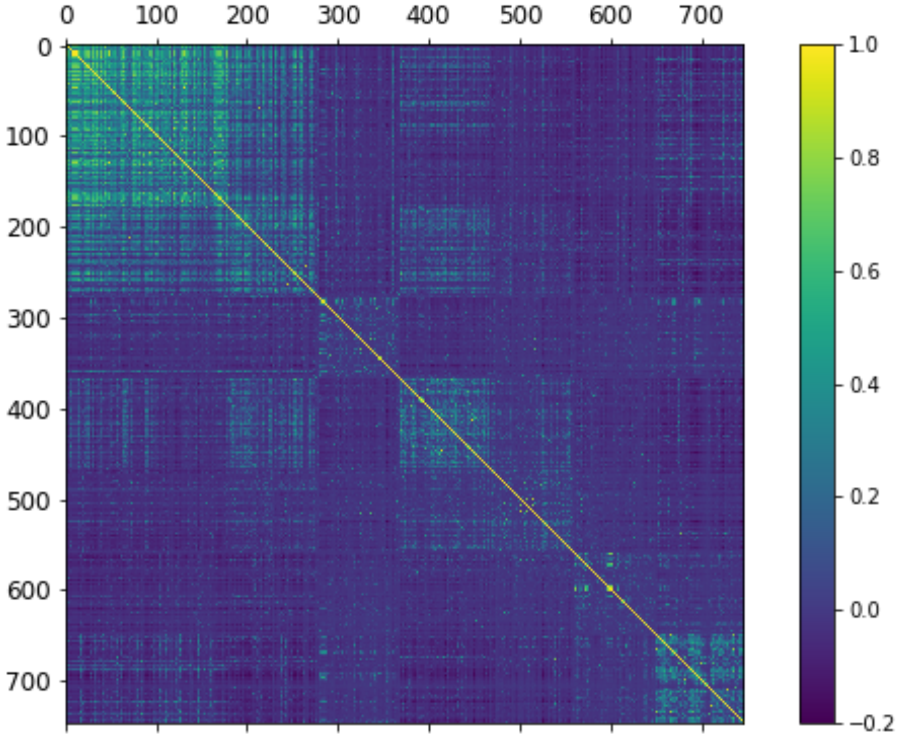


**Figure S5.** ***K*-mean clustering cross-correlation matrix (*k* = 7 model order).** The experiment (*e*) x experiment (*e*) cross-correlation matrix ordered by the seven-MAG solution from the *k* = 7 model order indicated that between-group differences of experiment correlation distributions were maximized while within-group differences were minimized.

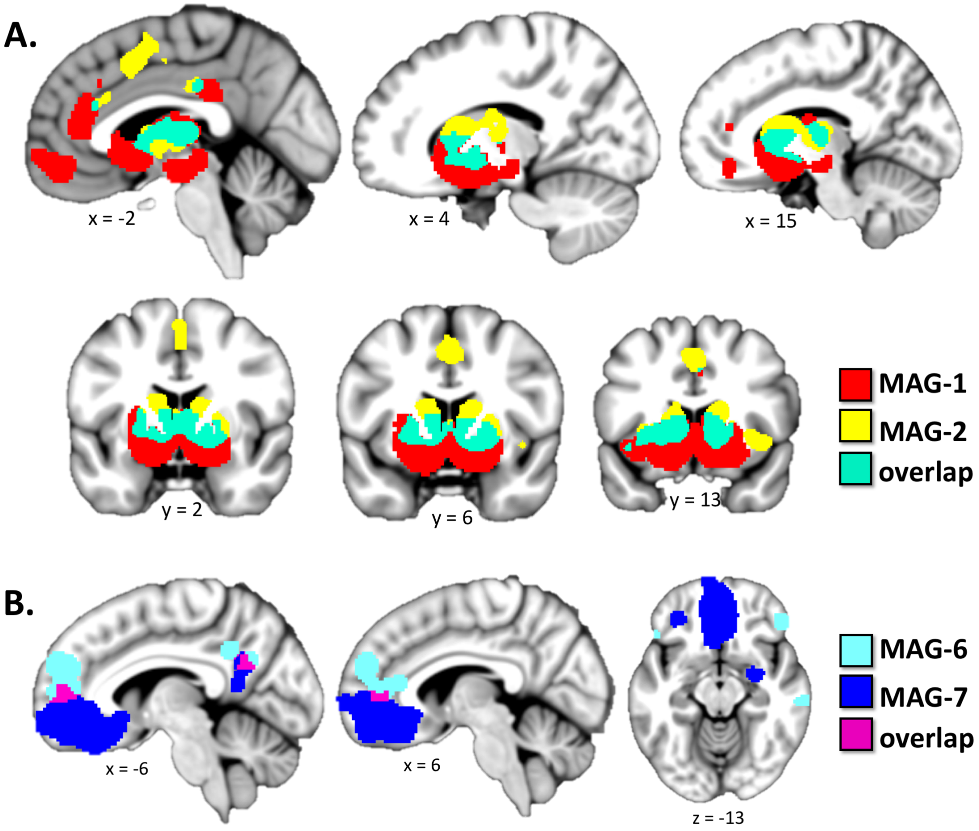


<u>Figure S6.</u> **Comparison of convergent activity for MAG-1 vs. -2 and for MAG-6 vs. -7.** (**A**) Whereas, both MAG-1 (red) and MAG-2 (yellow) displayed convergent activity that overlapped (cyan) in the striatum and medial frontal cortex, MAG-1's convergent activity was located more ventrally. Overlapping activity of these MAGs (cyan) was observed in the mid-striatum while activity unique to MAG-1 (red) was localized to the ventral striatum and ventromedial prefrontal cortex and the activity unique to MAG-2 (yellow) was localized to the dorsal striatum and dorsal medial frontal cortex. (**B**) MAG-6 (aqua) displayed convergent activity in the central medial prefrontal cortex and the precuneus, whereas MAG-7 (dark blue) displayed convergent activity located just ventral to that of MAG-6 in the ventral orbital frontal cortex and posterior cingulate. There was modest overlap of these MAGs in the medial prefrontal cortex and posterior cingulate (magenta). Additionally, MAG-6 displayed clusters of convergent activity in the lateral inferior prefrontal gyrus, middle temporal gyrus, and temporal parietal junction, whereas MAG-7 displayed convergent activity in the right amygdala.

### **MANUAL (vs. automated) ANNOTATION OF EXPERIMENTS FOR FUNCTIONAL DECODING**

**Corpus-specific manual annotation of experiments (*k* = 5 solution): Rationale.** The manual annotation of our corpus was prompted by our observation that the existing annotation of experimental contrasts provided by the BrainMap taxonomy was too generalized and nonspecific to capture the nuanced, yet critical distinctions of the specific reward-related neuroimaging contrasts. Thus, our functional decoding required metadata terms that were capable of capturing the precise nature of each experimental contrast. To achieve this level of detail, we performed corpus-specific, manual annotations that relied on the generation of an experiment-specific glossary through an admittedly subjective process. We acknowledge that this technique could be improved and have thus decided to instead perform and present a more automated and objective (yet, also more general) functional decoding technique in the main text utilizing a NeuroSynth-based strategy. However, as our manual decoding approach provides an example of the initial steps taken towards developing corpus- and domain-specific (e.g., reward processing) annotation ontologies, we have included methodological details and results from the *k* = 5 solution below.

To more precisely and succinctly characterize mental operations associated with each experimental contrast, we coded each one with cognitive-behavioral terms through a blind, multi-rater procedure. Due to the nuanced, yet critical distinctions between specific neuroimaging contrasts included in our reward processing corpus of results, our functional decoding required metadata terms capable of capturing the precise nature of each experimental contrast. To achieve this level of detail, we performed corpus-specific, manual annotations using a newly generated glossary of terms reflecting a summative definition of commonly operationalized phenomena in the included reward processing papers. First, a glossary was created consisting of terms (**Table**

**S6**) commonly employed to describe cognitive-behavioral aspects of reward processing tasks in the corpus. To create this glossary, raters read each published article's method section and collapsed synonyms, used to describe similar task contexts, task events, and/or cognitive phenomena into a singular term that encompassed a summative meaning. This resulted in a glossary of 42 terms which served to reduce potential, unnecessary lexical variability while still providing needed specificity to more fully capture the multifaceted aspects of reward processing tasks. A brief definition of each glossary term can be found in **Table S6**. Then to annotate all experimental contrasts, these glossary terms were assigned to each experimental contrast based on a review of the original article and the associated BrainMap metadata. Discrepancies between raters were discussed until consensus was reached. The "experiment name" BrainMap metadata was taken into particular consideration during this annotation process as it often provided the most specific definition of the contrast. Each contrast could be coded with multiple terms if all the terms appropriately pertained to the contrast.

**<u>Table S6.</u> Corpus-specific manual annotations: Term glossary and frequency distribution across MAGs.**

| Term | Definition | Total Frequency | Frequency for MAG-1 | Frequency for MAG-2 | Frequency for MAG-3 | Frequency for MAG-4 | Frequency for MAG-5 |
|---|---|---|---|---|---|---|---|
| **MID** | contrast isolated part of a monetary incentive delay task | 199 | 74 | 35 | 31 | 26 | 33 |
| **positive outcomes** | contrast represented an instance in which a positive outcome was delivered | 163 | 63 | 27 | 20 | 18 | 35 |
| **gambling choice** | contrast represented an instance in which the participant chooses between two or more outcome contingencies that varied in either risky-ness, probability or another, similar parameter | 132 | 36 | 19 | 27 | 37 | 13 |
| **value** | contrast isolated the value of an outcome, for example: a contrast that subtracted situations in which the outcome was $5 from situations in which the outcome was $1 | 102 | 27 | 15 | 16 | 20 | 24 |
| **negative outcomes** | contrast represented an instance in which a negative outcome was delivered | 91 | 13 | 23 | 19 | 27 | 9 |
| **prediction uncertainty** | contrast represented an instance in which participants had to choose between options that would lead to different outcomes without being completely sure which option led to which outcome | 83 | 8 | 8 | 13 | 36 | 18 |
| **choice** | contrast represented an instance in which a participant made a choice | 79 | 18 | 14 | 10 | 20 | 17 |
| **anticipation** | contrast isolated the period before an expected outcome was delivered | 73 | 28 | 14 | 11 | 10 | 10 |
| **reward learning** | contrast represented an algorithm that calculated how task performance changed over time | 68 | 10 | 5 | 12 | 26 | 15 |
| **delay** | contrast represented a wait period (real or hypothetical) before an outcome included in delay discounting tasks | 62 | 10 | 8 | 15 | 22 | 7 |
| **reversal learning task** | contrast isolated a part of any task in which outcome contingencies were periodically changed throughout the task | 59 | 5 | 5 | 14 | 23 | 12 |
| **reward price** | contrast represented either how much a participant would pay for an outcome or the delivery of information about the cost of an outcome | 59 | 8 | 5 | 12 | 12 | 22 |
| **delay discounting** | contrast represented part of a task that involved making decisions based on the delay until an outcome | 57 | 10 | 7 | 15 | 17 | 8 |
| **rewarded performance** | contrast represented an instance in which any positive outcome, that was contingent on a task response, was delivered | 54 | 9 | 15 | 8 | 14 | 8 |
| **probability** | contrast represented the probability of an outcome | 53 | 12 | 7 | 12 | 17 | 5 |
| **reward omission** | contrast represented instances in which an expected positive outcome was not delivered | 52 | 26 | 9 | 2 | 11 | 4 |
| **risk** | contrast represented the odds of an outcome contingency | 51 | 10 | 12 | 3 | 18 | 8 |
| **social** | contrast represented part of a task that involved interacting with another person (real or hypothetical) | 42 | 9 | 10 | 9 | 7 | 7 |

| | | | | | | | |
|---|---|---|---|---|---|---|---|
| **wheel of fortune task** | contrast isolated part of a task that involved a 'wheel of fortune.' Often the participant was instructed to choose a section of the wheel and that section would indicate the outcome delivered if the wheel landed on that section when spun. | 42 | 12 | 5 | 11 | 10 | 4 |
| **negative outcomes escape** | contrast represented an instance in which a negative outcome was avoided | 37 | 8 | 4 | 5 | 10 | 10 |
| **food** | contrast isolated part of a task that involved food (real or hypothetical) | 35 | 10 | 8 | 9 | 2 | 6 |
| **picture of desired object** | contrast isolated part of a task that involved a picture of a desired object | 33 | 6 | 1 | 7 | 10 | 9 |
| **probabilistic conditioning** | contrast isolated part of a task that involved a probabilistic stimuli-outcome contingency | 31 | 9 | 3 | 5 | 4 | 10 |
| **classical conditioning** | contrast isolated part of a task that involved stimuli-outcome associations | 27 | 14 | 4 | 3 | 3 | 3 |
| **performance feedback** | contrast isolated an instance in which feedback about task performance was provided | 26 | 6 | 7 | 3 | 5 | 5 |
| **risk vs. amount** | contrast isolated part of a task in which participants choose between outcomes varying in risk and value. In most cases, the choice was between a high-risk, high-value outcome, and a low-risk, low-value outcome | 23 | 1 | 6 | 5 | 9 | 2 |
| **go/no-go** | contrast isolated part of a go/no-go inhibitory control task | 17 | 4 | 3 | 4 | 3 | 3 |
| **moving through virtual maze** | contrast isolated part of a task in which participants had to navigate through a virtual space | 17 | 2 | 4 | 1 | 1 | 9 |
| **predator and prey paradigm** | contrast isolated part of the predator and prey paradigm in which participants had to navigate a virtual space to escape a 'predator.' | 17 | 2 | 4 | 1 | 1 | 9 |
| **unexpected outcomes** | contrast isolated the delivery of unexpected outcomes | 15 | 6 | 4 | 3 | 2 | 0 |
| **pattern recognition** | contrast isolated part of a task that required participants to learn associations between outcomes and complex patterns of stimuli or cues | 14 | 0 | 8 | 3 | 3 | 0 |
| **purchasing** | contrast isolated part of a task that involved paying a cost of some sort, to receive an outcome | 14 | 2 | 0 | 6 | 2 | 4 |
| **slot machine** | contrast represented part of a task involving a simulated slot machine | 13 | 3 | 3 | 0 | 5 | 2 |
| **stock market** | contrast represented part of a task involving the monetary decisions and outcomes of a simulated stock market | 12 | 1 | 3 | 4 | 3 | 1 |
| **tower of London task** | contrast represented part of a Tower of London task. These tasks involve cognitive control, planning and problem solving | 9 | 1 | 3 | 0 | 1 | 4 |
| **valence** | contrast isolated the difference between positive and negative outcomes | 7 | 3 | 0 | 2 | 1 | 1 |
| **altruistic donation** | contrast represented part of a task that involved making decisions about voluntarily giving valuable capital to others | 6 | 3 | 0 | 1 | 1 | 1 |
| **face attractiveness** | contrast represented part of a task instructing participants to make | 6 | 1 | 0 | 0 | 1 | 4 |

| | | | | | | | |
|---|---|---|---|---|---|---|---|
| | judgments and indicate preferences about human faces | | | | | | |
| **advice** | contrast represented part of a task in which a participant had to make a choice and were given advice from an outside source (usually an 'expert') on which choice to make. | 3 | 2 | 0 | 0 | 1 | 0 |
| **reward delivery** | contrast isolated an instance in which a positive outcome was delivered | 3 | 1 | 0 | 0 | 2 | 0 |
| **blackjack** | contrast represented part of a task in which participants played a type of blackjack game | 2 | 1 | 1 | 0 | 0 | 0 |
| **verbal reward** | contrast isolated an instance in which positive verbal (auditory or written) feedback was provided | 2 | 1 | 1 | 0 | 0 | 0 |

Note. Terms used to code all experimental contrasts in corpus with a brief definition describing rational for assigning it to a contrast. Each term's frequency across the corpus and frequency for each MAG are provided.

**Corpus-specific manual annotations for functional decoding of meta-analytic groupings (*k* = 5 solution): Methods.** To generate cognitive-behavior profiles for each MAG, we performed exploratory functional decoding analyses using the terms coding each experimental contrast. We characterized term frequency distributions within and across MAGs using an adapted implementation of the forward and reverse inference analyses employed by NeuroSynth to calculate term frequency distributions within and across voxels from a large pool of studies (Yarkoni, Poldrack, Nichols, Van Essen, & Wager, 2011). Forward inference analyses have been used to characterize the likelihood of activation given a mental phenomenon, and reverse inference analyses have been used to characterize the likelihood of a mental phenomenon given activation (Cieslik et al., 2013; Nickl-Jockschat et al., 2015; Poldrack, 2006; Yarkoni et al., 2011). In our adaptation of these analyses, multiple comparisons across terms were corrected for using the Benjamini-Hochberg procedure, which restricts the false discovery rate (FDR) to a given level (0.05; (Benjamini & Hochberg, 1995).

We first performed a within-MAG consistency analysis which calculated the probability of a term given assignment to a particular MAG, P(term | MAG). This analysis identified terms with a higher representation in a certain MAG than would be expected given a baseline level, which was defined as that MAG's average term frequency. Significance was assessed with one-way Chi-square tests of independence (FDR-corrected for the number of terms [$p_{FDR\text{-}corrected}<0.05$]). A significant, positive association between a term and a MAG indicated that the term was coded for experiments in that particular MAG at a higher frequency than would be expected given the average frequency of all terms coded for that MAG. An across-MAG selectivity analysis was then performed which calculated the probability of experiment MAG assignment given a term, P(MAG | term). This analysis assessed whether a term had a higher frequency within a certain MAG than would be expected given the frequency of that term across all other MAGs. Significance was assessed with two-way Chi-square tests ($p_{FDR\text{-}corrected}$<0.05) with a significant, positive association between a term and a MAG indicating that the term was coded for experiments in that particular MAG at a higher frequency than would be expected given the frequency in all other MAGs. Whereas the within-MAG analysis was influenced by each term's overall frequency in the corpus but not by cluster size (i.e., number of experimental contrasts in a MAG), the across-MAG analysis was less influenced by term frequency but more influenced by cluster size. Consequently, both the within-MAG and across-MAG analyses provided complementary information about term-MAG associations and were both utilized to create behavior profiles for each MAG. The posterior probability of each association was calculated (assuming a uniform prior of 0.5) as a measure of effect size (Poldrack, 2006), and plotted for significant terms from each MAG (**Fig. S8**).

**Corpus-specific manual annotations for functional decoding of meta-analytic groupings (*k* = 5 solution): Results.** We calculated the frequency at which each term was coded across the entire corpus, in addition to the frequency it was coded in each MAG (**Table S6**). We then employed within-MAG consistency and across-MAG selectivity analyses that utilized these term frequency distributions to determine statistically significant term-MAG associations. We use the phrase 'behavior profile' to refer to the collection of meta-analytic terms/labels showing a significant, positive association with a MAG from either the within-MAG consistency or across-MAG selectivity analyses ($p < 0.05$). If a term was significantly associated with at least four of the five MAGs, it was assigned to a 'common reward processing behavior profile'. This common behavior profile (**Fig. S7**) was composed of five terms: *Monetary Incentive Delay task* (MID) and *positive outcomes* (associated with all 5 MAGs) as well as *choice*, *value,* and *gambling choice* (associated with 4 out of the 5 MAGs). All other terms (not included in the common behavior profile) that displayed a significant association with a MAG ($p < 0.05$) were included in that MAG's unique behavior profile. We found that results from this corpus-specific, manual-annotation decoding procedure largely supported functional decoding results for the $k = 5$ solution that were derived using a NeuroSynth strategy (**Fig. S2**).

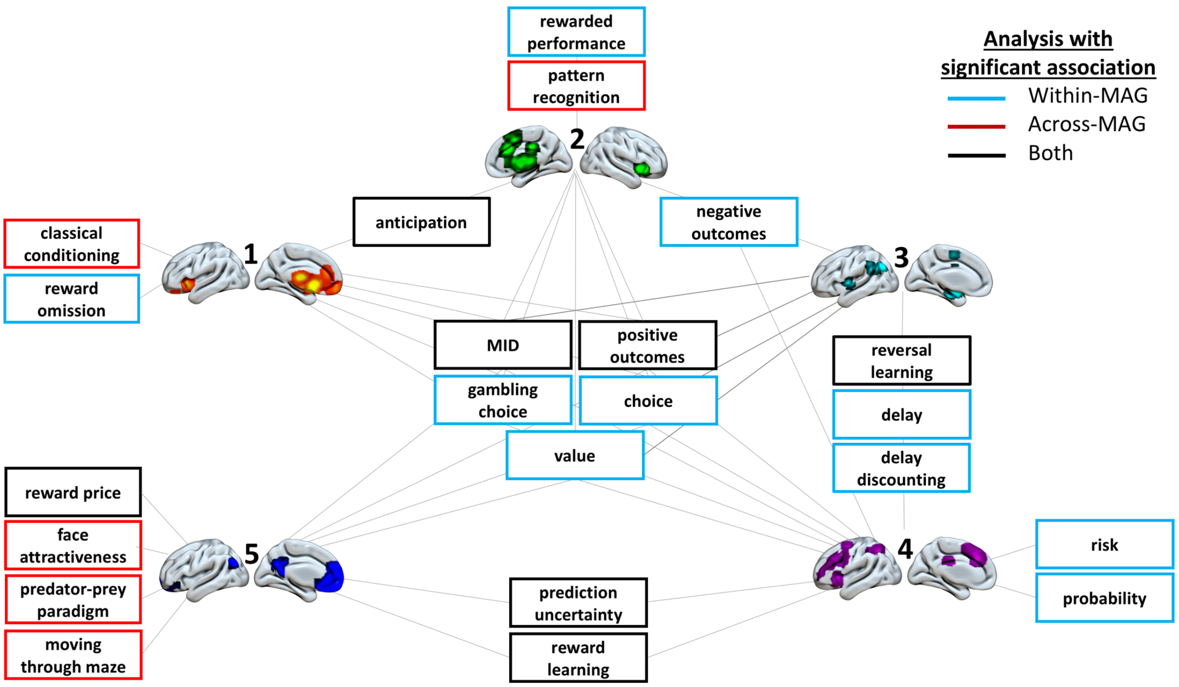


**Figure S7. Behavior profiles associated with each meta-analytic grouping (MAG) of reward processing experiments utilizing corpus-specific manual annotations ($k = 5$ solution).** Behavior profiles for each MAG consisted of terms demonstrating significant positive associations in either the within-MAG consistency analysis (blue outline), the between-MAG selectivity analysis (red), or both analyses (black) ($p_{FDR\text{-}corrected} < 0.05$). Terms significantly associated with at least four of the five MAGs were assigned to the 'common reward processing behavior profile'. Terms significantly and positively associated with only one MAG were included in that MAG's unique behavior profile.

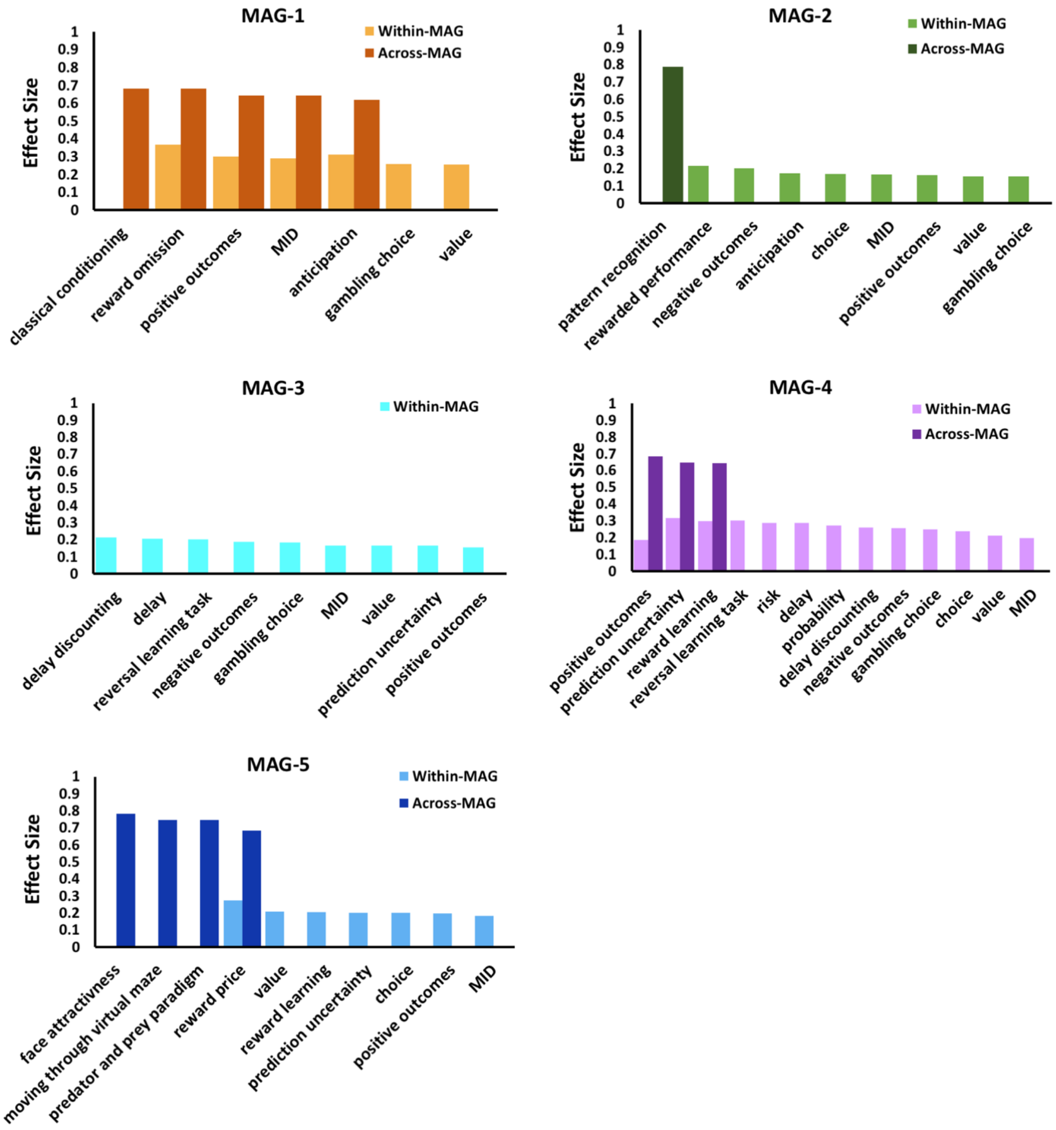


**<u>Figure S8.</u> Effect size of each significant term-MAG association identified using corpus-specific manual annotations (*k* = 5 solution)**. Posterior probabilities were calculated and used as a measure of effect size for terms (out of a possible 42 terms) that were identified as significant in the within-MAG (lighter colors) or across-MAG functional decoding analysis (darker colors). The within-MAG analysis effect size indicates the probability of term, given a MAG (P[term | MAG]), while the across-MAG analysis effect size indicates the probably of a MAG, given a term (P[MAG | term]). The posterior probability of each association was calculated (assuming a uniform prior of 0.5), used as a measure of effect size (Poldrack, 2006), and plotted for significant terms from each MAG. The uniform prior was employed primarily to prevent estimates of posterior probabilities from being overwhelmed by differences in terms' base frequencies across the reward processing literature. In all instances the term-MAG associations reaching significance in the across-MAG analysis had higher effect sizes than term-MAG associations reaching significance in the within-MAG analysis. Additionally, terms uniquely associated with a certain MAG usually had higher effect sizes than terms significantly associated with multiple MAGs. Examining the effect sizes of significant terms increases the transparency of the functional decoding analysis.

### SUPPELMENTAL REFERENCES

Benjamini, Y., & Hochberg, Y. (1995). Controlling the False Discovery Rate: A Practical and Powerful Approach to Multiple Testing. *Journal of the Royal Statistical Society, 57*(1), 289-300.

Cieslik, E. C., Zilles, K., Caspers, S., Roski, C., Kellermann, T. S., Jakobs, O., . . . Eickhoff, S. B. (2013). Is there "one" DLPFC in cognitive action control? Evidence for heterogeneity from co-activation-based parcellation. *Cereb Cortex, 23*(11), 2677-2689. doi:10.1093/cercor/bhs256

Nickl-Jockschat, T., Rottschy, C., Thommes, J., Schneider, F., Laird, A. R., Fox, P. T., & Eickhoff, S. B. (2015). Neural networks related to dysfunctional face processing in autism spectrum disorder. *Brain Struct Funct, 220*(4), 2355-2371. doi:10.1007/s00429-014-0791-z

Poldrack, R. A. (2006). Can cognitive processes be inferred from neuroimaging data? *Trends Cogn Sci, 10*(2), 59-63. doi:10.1016/j.tics.2005.12.004

Yarkoni, T., Poldrack, R. A., Nichols, T. E., Van Essen, D. C., & Wager, T. D. (2011). Large-scale automated synthesis of human functional neuroimaging data. *Nature methods, 8*(8), 665.

Benjamini, Y., & Hochberg, Y. (1995). Controlling the False Discovery Rate: A Practical and Powerful Approach to Multiple Testing. *Journal of the Royal Statistical Society, 57*(1), 289-300.

Cieslik, E. C., Zilles, K., Caspers, S., Roski, C., Kellermann, T. S., Jakobs, O., . . . Eickhoff, S. B. (2013). Is there "one" DLPFC in cognitive action control? Evidence for heterogeneity from co-activation-based parcellation. *Cereb Cortex, 23*(11), 2677-2689. doi:10.1093/cercor/bhs256

Nickl-Jockschat, T., Rottschy, C., Thommes, J., Schneider, F., Laird, A. R., Fox, P. T., & Eickhoff, S. B. (2015). Neural networks related to dysfunctional face processing in autism spectrum disorder. *Brain Struct Funct, 220*(4), 2355-2371. doi:10.1007/s00429-014-0791-z

Poldrack, R. A. (2006). Can cognitive processes be inferred from neuroimaging data? *Trends Cogn Sci, 10*(2), 59-63. doi:10.1016/j.tics.2005.12.004

Yarkoni, T., Poldrack, R. A., Nichols, T. E., Van Essen, D. C., & Wager, T. D. (2011). Large-scale automated synthesis of human functional neuroimaging data. *Nature methods, 8*(8), 665.

Benjamini, Y., & Hochberg, Y. (1995). Controlling the False Discovery Rate: A Practical and Powerful Approach to Multiple Testing. *Journal of the Royal Statistical Society, 57*(1), 289-300.

Cieslik, E. C., Zilles, K., Caspers, S., Roski, C., Kellermann, T. S., Jakobs, O., . . . Eickhoff, S. B. (2013). Is there "one" DLPFC in cognitive action control? Evidence for heterogeneity from co-activation-based parcellation. *Cereb Cortex, 23*(11), 2677-2689. doi:10.1093/cercor/bhs256

Nickl-Jockschat, T., Rottschy, C., Thommes, J., Schneider, F., Laird, A. R., Fox, P. T., & Eickhoff, S. B. (2015). Neural networks related to dysfunctional face processing in autism spectrum disorder. *Brain Struct Funct, 220*(4), 2355-2371. doi:10.1007/s00429-014-0791-z

Poldrack, R. A. (2006). Can cognitive processes be inferred from neuroimaging data? *Trends Cogn Sci, 10*(2), 59-63. doi:10.1016/j.tics.2005.12.004

Yarkoni, T., Poldrack, R. A., Nichols, T. E., Van Essen, D. C., & Wager, T. D. (2011). Large-scale automated synthesis of human functional neuroimaging data. *Nature methods, 8*(8), 665.
